# Pre-symptomatic proteomic and metabolomic profiling identifies compensated ER-redox-metabolic adaptation and early nuclear vulnerability in neuronal ERO1L toxicity

**DOI:** 10.64898/2026.09.06.749746

**Authors:** Luca Lo Piccolo, Ranchana Yeewa, Puttachat Poound, Chansunee Panto, Jutarop Phetcharaburanin, Manida Suksawat, Jakkapong Kluebsoongnoen, Wassana Jamnongkarn, Pipob Suwanchaikasem, Salinee Jantrapirom

**Affiliations:** Center of Multidisciplinary Technology for Advanced Medicine (CMUTEAM), Faculty of Medicine, Chiang Mai University, Chiang Mai, Thailand; Drosophila Centre for Human Diseases and Drug Discovery (DHD), Faculty of Medicine, Chiang Mai University, Chiang Mai, Thailand; Department of Systems Biosciences and Computational Medicine, Faculty of Medicine, Khon Kaen University, Khon Kaen, Thailand; The National Phenome Institute, Office of the President, Khon Kaen University, Khon Kaen, Thailand; Baiya Phytopharm Co., Ltd., Bangkok 10330, Thailand; Department of Pharmacology, Faculty of Medicine, Chiang Mai University, Chiang Mai, Thailand

**Keywords:** ERO1L, brain aging, multi-omics, ER-redox homeostasis, chromatin stability, *Drosophila melanogaster*

## Abstract

Aging progressively challenges neuronal proteostasis, redox homeostasis, and metabolism, yet the molecular changes that precede functional decline remain poorly understood. Endoplasmic reticulum oxidoreductin 1 (ERO1), a key regulator of oxidative protein folding, links endoplasmic reticulum (ER) proteostasis with cellular redox balance and is elevated in aging and neurodegenerative contexts. Here, we investigated how neuronal ERO1L elevation reshapes cellular homeostasis before overt dysfunction in *Drosophila melanogaster*. Endogenous *ERO1L* expression increased with age, and neuronal *ERO1L* elevation shortened lifespan and caused progressive locomotor decline. This effect was strongly cell-type dependent, as *ERO1L* elevation in glia, muscle, or fat body did not produce a comparable survival phenotype. At day 5 post-eclosion, locomotor performance remained preserved and major brain reactive-oxygen-species (ROS) accumulation was not yet detectable, defining a pre-symptomatic stage. Multi-omic profiling at this stage revealed selective remodelling of ER proteostasis, redox defence, and mitochondrial-energy pathways, together with changes in central-carbon, nitrogen, and purine metabolism. In contrast to these broadly adaptive responses, chromatin- and RNA-homeostasis-associated proteins were selectively reduced, accompanied by decreased *HP1*, *dFmr1*, and *Piwi* expression and increased transposable-element transcripts. Thus, neuronal ERO1L elevation establishes a pre-symptomatic state in which proteostatic and metabolic adaptation coexists with early vulnerability of nuclear and RNA-homeostasis pathways, preceding overt oxidative stress and behavioural decline. These findings provide an in vivo framework to investigate how age-associated *ERO1L* elevation may progressively reduce neuronal resilience during brain aging.

## Introduction

Aging progressively challenges neuronal proteostasis, redox balance and energy metabolism (Coleman et al., 2026). Neurons are particularly susceptible because they are long-lived, highly polarized and metabolically demanding, requiring sustained protein quality control and organelle homeostasis throughout life. Disturbances in these systems are prominent features of brain aging and neurodegenerative disease (NDs), yet functional decline often becomes apparent only after molecular and cellular responses have been engaged for an extended period (Grochowska et al., 2022; Hipp et al., 2019). Defining the early states that preserve neuronal function is therefore important for distinguishing adaptive responses from vulnerabilities that may precede and contribute to degeneration.

Endoplasmic reticulum oxidoreductin 1 (ERO1) is positioned at the intersection of proteostasis, redox regulation and cellular metabolism. ERO1 is a conserved ER-resident, flavin adenine dinucleotide (FAD)-dependent oxidase that reoxidizes protein disulfide isomerases (PDIs), thereby sustaining disulfide-bond formation in secretory and membrane proteins (Shergalis et al., 2020). Electron transfer through the ERO1–PDI relay ultimately reduces molecular oxygen to hydrogen peroxide (H_2_O_2_), coupling ER protein-folding demand to local redox balance (Ramming et al., 2015; Wang and Wang, 2023). ERO1 activity is tightly regulated through reversible disulfide bonds and interactions with PDI-family proteins, allowing oxidative folding to adapt to changing redox conditions (Benham et al., 2013; Inaba et al., 2010). In mammals, two ERO1 paralogues are present: ERO1A, the broadly expressed isoform also known as ERO1L, and the more tissue-restricted ERO1B (Pagani et al., 2000). ERO1A is also induced by ER stress and hypoxia-responsive programmes, and has been linked to ER calcium release, ER-mitochondria communication and mitochondrial bioenergetics (Bassot et al., 2023; Gilady et al., 2010; Li et al., 2009; Renna et al., 2026; Varone et al., 2021). Altered ERO1A activity may therefore influence neuronal physiology through coordinated effects on protein folding, redox balance, calcium signalling and mitochondrial metabolism.

Consistent with this broader role, ERO1A is elevated in several NDs and stress-associated settings, where increased ERO1 activity has been associated with oxidative stress, calcium dysregulation and pro-apoptotic signalling (Jha et al., 2023; Lehtonen et al., 2016; Maguire et al., 2025; Qiao et al., 2022; Yeewa et al., 2024; Zhang et al., 2022). Notably, ERO1A expression is also increased during human cellular senescence, as reported in the Senescence-Associated Secretory Phenotype (SASP) Atlas (Basisty et al., 2020), providing evidence that regulation of this pathway is associated with the aging process itself. ER-stress responses are likewise observed across models of Alzheimer’s disease, Parkinson’s disease, amyotrophic lateral sclerosis (ALS) and other disorders characterized by proteostasis disruption. However, the consequences of ERO1 induction are highly context dependent, varying with stress intensity and duration, cellular metabolic state and disease stage. Increased ERO1A in aging and disease may therefore represent an adaptive response to increased protein-folding demand, a source of additional redox stress, or a transition between these states.

Animal studies support a functional, dose- and time-dependent role for ERO1 in stress resilience and lifespan. In *Caenorhabditis elegans*, adult-stage RNA interference against the ERO1 orthologue *ero-1* extends lifespan, whereas early depletion disrupts development (Curran and Ruvkun, 2007). In *Drosophila melanogaster*, Ero1L is the orthologue of human ERO1A (hereafter refered as to ERO1L) (Yeewa et al., 2024). Reducing ERO1L activity mitigates neuronal ER stress, improves disease-associated phenotypes and extends lifespan under environmental or pathological ER stress, whereas neuronal ERO1L elevation shortens lifespan during ER-stress challenge (Yeewa et al., 2024). These findings establish ERO1A as a modulator of neuronal stress tolerance and organismal survival, but leave unresolved how neurons accommodate sustained ERO1A elevation before oxidative injury and functional decline emerge. Defining this early state is important for distinguishing compensatory responses that preserve neuronal function from molecular vulnerabilities that may precede overt decline.

However, the molecular burden imposed by chronic neuronal ERO1A elevation therefore remains poorly defined. In particular, it is unclear how sustained ERO1A activity reshapes the interplay between ER protein folding, redox regulation and cellular metabolism, and which molecular responses emerge while neuronal function is still preserved. *Drosophila* provides an opportunity to address these questions through cell-type-specific genetic control, a tractable aging trajectory and established behavioural and molecular readouts. Here, we used pan-neuronal ERO1L elevation and combined longitudinal phenotypic characterization with proteomic, metabolomic and targeted molecular analyses to define the cellular consequences of sustained ERO1A activity and identify early molecular features associated with neuronal resilience.

## Methods

### *Drosophila* stocks and husbandry

*Drosophila melanogaster* stocks were maintained on standard cornmeal-yeast-glucose medium at 22 °C and 60% relative humidity under a 12 h:12 h light:dark cycle, unless otherwise indicated. The following lines were obtained from the Bloomington *Drosophila* Stock Center (BDSC): nSyb-GAL4 (BDSC #68222), yw (BDSC #1495), UAS-*Ero1L* (BDSC #42698), repo-GAL4 (BDSC #7415), Lsp2-GAL4 (BDSC #6357), Mhc-GAL4 (BDSC #55132), UAS-*LacZ* (BDSC #1776), and UAS-*ERO1L* RNAi (BDSC #64674). Tissue-specific *ERO1L* overexpression (ERO1L-OE) or RNAi-mediated knockdown (ERO1L-IR) was achieved using the GAL4/UAS binary system (Brand and Perrimon, 1993). Crosses were established at 25 °C, and all experiments were conducted at 25 °C unless otherwise stated. Adult males were collected within 24 h of eclosion and aged at a standardized density of 15 flies per vial. To reduce genetic-background variation, lines were independently backcrossed for six generations into the yw background.

### Lifespan assays

For lifespan analysis, virgin females carrying the relevant GAL4 driver were crossed with males carrying UAS-*ERO1L* on standard cornmeal-yeast-glucose medium. Crosses were established in parallel using 10 virgin females and 10 males per vial, with three independent parental crosses prepared for each genotype. Control crosses were generated in parallel using the corresponding non-driven UAS-*ERO1L* line. Newly eclosed male progeny were collected within 24 h of eclosion and randomly assigned to lifespan cohorts. For each genotype, 100 males were distributed across 10 independent vials at a density of 10 flies per vial. Flies were maintained on standard medium at 25 °C under the humidity and light:dark conditions described above. Living flies were transferred without anaesthesia to fresh food every 3 days, and deaths were recorded at each transfer until all flies had died. Flies lost through accidental death or escape were recorded as censored observations. All lifespan groups were established in parallel within the same experimental batch and maintained under identical conditions. Kaplan-Meier survival curves were generated from individual-fly survival data, and differences between genotypes were evaluated using the log-rank (Mantel-Cox) test. Standard lifespan workflows similarly use multiple replicate vials, regular food transfers with mortality scoring, censored observations, Kaplan-Meier curves, and log-rank testing.

### Negative-geotaxis locomotor assay

Age-dependent locomotor performance was assessed using a negative-geotaxis assay adapted from (Lo Piccolo et al., 2026a). Male flies were tested at 5, 10, 15, 20, and 25 days post-eclosion, with 100 age-matched males per genotype distributed across 10 independent vials (10 flies per vial). Flies were transferred without anesthesia to transparent conical tubes (15 cm height) and acclimatized for 10 min. Before each trial, flies were gently tapped to the bottom of the tube and their climbing behaviour was recorded for 5 s. Five consecutive trials were performed at 30-s intervals. Individual flies were scored according to the maximum height reached during the 5-s observation period: 0, <2.0 cm; 1, 2.0–3.9 cm; 2, 4.0–5.9 cm; 3, 6.0–7.9 cm; 4, 8.0–9.9 cm; and 5, ≥10 cm. A climbing index was calculated for each vial as the weighted mean score across all flies. The mean climbing index across the five trials was used for statistical analysis. Genotypes were tested in parallel with age-matched controls.

### RNA extraction, cDNA synthesis, and RT–qPCR

Total RNA was isolated from adult *Drosophila* heads using the RNeasy Mini Kit (QIAGEN) according to the manufacturer’s instructions. For longitudinal *ERO1L* expression analysis, RNA was extracted from pools of 50 heads per biological replicate, with three independent biological replicates analysed per age group. For transposable-element (TE) expression analysis, RNA was extracted from pools of 50 heads per biological replicate, with four independent biological replicates per genotype. RNA concentration and purity were assessed using a NanoDrop One/OneC spectrophotometer (Thermo Fisher Scientific). For cDNA synthesis, 1 μg of total RNA was reverse transcribed using the SensiFAST cDNA Synthesis Kit (Bioline). Quantitative RT–PCR was performed in technical triplicate using SensiFAST SYBR Lo-ROX (Bioline) on a CFX Opus Real-Time PCR System (Bio-Rad). Primer sequences are provided in Supplementary Table 1. Relative *ERO1L* transcript abundance was normalized to *Act5C*, whereas TE transcript abundance was normalized to *GAL4*. Relative expression was calculated using the 2^−ΔΔCt method. For longitudinal *Ero1L* analysis, expression values were normalized to the designated reference age group. For TE analysis, relative transcript abundance in ERO1L-OE heads was normalized to the corresponding LacZ control group.

### Dihydroethidium staining and quantification of brain fluorescence

The Dihydroethidium (DHE) staining protocol was adapted from a previously described method (Yeewa et al., 2026) with modifications. Brains from adult flies at 5 or 10 days post-eclosion were dissected in ice-cold Schneider’s insect medium (PAN Biotech, P04-91500, Germany) and immediately incubated in 1× PBS containing 30 µM dihydroethidium (DHE) (Supplementary Table 2) for 5 min at 25 °C in the dark. Following staining, brains were fixed in 4% paraformaldehyde (PFA) at 25 °C for 5 min, washed three times in 1× PBS, and mounted on glass slides using spacers and VECTASHIELD antifade mounting medium. Samples were imaged within 10 min of mounting using an Olympus FV3000 confocal laser-scanning microscope (Olympus, Japan). DHE fluorescence was quantified using Olympus cellSens Imaging Software. Brains from all experimental groups within each experiment were processed, mounted, and imaged in parallel using identical microscope and acquisition settings. A standardized region of interest (ROI) encompassing the brain region of interest was applied to each sample. Background fluorescence was measured in a cell-free area of the corresponding image and subtracted from the ROI fluorescence signal. Each brain was considered one biological replicate for statistical analysis.

### Whole-mount brain immunofluorescence

Brains from adult flies at 5 days post-eclosion were dissected in ice-cold 1× PBS and maintained on ice throughout the dissection, as previously described in (Lo Piccolo et al., 2026b), with few modifications. Briefly, brains were fixed in 4% PFA in PBS for 30 min at room temperature (RT) with gentle rocking, followed by three washes in PBS containing 0.3% Triton X-100 (PBST) for 20 min each at RT. Samples were blocked for 1 h at RT in PBST containing 5% normal goat serum (NGS). Brains were incubated for 48 h at 4 °C with mouse anti-HP1 antibody (clone C1A9; Developmental Studies Hybridoma Bank, DSHB; 1:50) diluted in blocking buffer with gentle rotation. Following three washes in PBST for 20 min each at RT, brains were incubated overnight at 4 °C in the dark with goat anti-mouse Alexa Fluor 594-conjugated secondary antibody (1:500) and Alexa Fluor 488-conjugated phalloidin (1:200), diluted in blocking buffer. Samples were then washed three times in PBST for 20 min each at RT and counterstained with DAPI (1 µg/mL) for 15 min at RT in the dark. Following two washes in PBS for 10 min each, brains were mounted in antifade mounting medium. Confocal images were acquired using 405-, 488-, and 561-nm excitation for DAPI, Alexa Fluor 488, and Alexa Fluor 594, respectively. All experimental groups within each experiment were processed and imaged using identical acquisition settings. Antibodies are listed in Supplementary Table 2.

### Quantification of HP1 immunofluorescence

HP1 immunoreactivity was quantified from confocal images acquired under identical imaging conditions. Nuclei were identified based on DAPI fluorescence, and 50 nuclei were analyzed per brain. For each nucleus, area, mean fluorescence intensity, and integrated density of the HP1 channel were measured using Fiji/ImageJ. Background fluorescence was measured in a cell-free region of the corresponding image. Corrected total cell fluorescence (CTCF) was calculated for each nucleus as CTCF = integrated density − (area × mean background fluorescence). CTCF values from the 50 nuclei within each brain were averaged to generate a single value for each biological replicate. Ten brains were analyzed per genotype (*n* = 10 biological replicates per genotype). Statistical analyses were performed using the mean CTCF value for each brain, with individual nuclei treated as technical subsamples rather than independent biological replicates.

### Immunoblotting

Adult *Drosophila melanogaster* flies at 5 days post-eclosion were frozen in liquid nitrogen, and heads were separated by vigorous vortexing for 45 s. Crude head lysates were prepared as previously described, with minor modifications. Briefly, 20 heads were homogenized in 100 µL of 4× SDS sample loading buffer (Merck) containing 1% 2-mercaptoethanol. At least six independent biological extracts were prepared per genotype. Lysates were centrifuged at 12,000 × *g* for 15 min at 4 °C, and 10 µL of cleared lysate was separated by 10% SDS-PAGE and transferred to Immobilon-P PVDF membranes (Merck) using a Trans-Blot Turbo system (Bio-Rad). Membranes were blocked with 5% skim milk in TBST and incubated overnight with primary antibodies diluted in blocking buffer: rabbit anti-Piwi (1:500), mouse anti-dFmr1 (clone 5A11; 1:1,000), mouse anti-Elav (1:1,000), mouse anti-HP1 (clone C1A9; 1:1,000), rabbit anti-histone H3 (clone 3H1; 1:1,000), rabbit anti-H3K9me2 (1:2,000), and mouse anti-H3K9me3 (1:5,000). After washing, membranes were incubated with HRP-conjugated secondary antibodies (1:3,000). Chemiluminescence was detected using WesternSure reagents (LI-COR) and a C-DiGit Blot Scanner (LI-COR). Band intensities were quantified and normalized to the corresponding loading control. Antibody sources and catalogue information are provided in Supplementary Table S2.

### Proteomic sample preparation and digestion

For label-free quantitative proteomic analysis, *Drosophila* head samples from the indicated experimental groups were extracted in 6 M urea in 50 mM Tris-HCl at a ratio of 3 mg tissue per 100 µL lysis buffer. Samples were homogenized with two metal beads using a tissue homogenizer (SWE-FP, ServiceBio) at 50 Hz for two 3-min cycles and centrifuged at 14,000 rpm for 10 min. Proteins were precipitated by addition of four volumes of ice-cold methanol, followed by centrifugation at 14,000 rpm for 10 min. Protein pellets were resuspended in 50 mM ammonium bicarbonate (ABC) and normalized to 0.49 mg/mL, corresponding to the lowest protein concentration among samples. An aliquot of 40 µL was reduced with 10 mM dithiothreitol (DTT) at 65 °C for 30 min and subsequently alkylated with 25 mM iodoacetamide (IAA) for 20 min at room temperature in the dark. Proteins were digested with 2 µL of trypsin (0.1 mg/mL) at 37 °C overnight. Digestion was terminated by addition of 10% formic acid (FA), and samples were centrifuged at 14,000 rpm for 10 min. The resulting supernatants were transferred to LC–MS vials for analysis.

### LC-MS/MS analysis

Peptides were analyzed by liquid chromatography-tandem mass spectrometry (LC-MS/MS) using an Agilent 6545XT LC-QTOF system equipped with an Agilent Peptide Map column (2.1 × 150 mm, 2.7 µm) maintained at 60 °C. A 20-µL aliquot of each sample was injected. Chromatographic separation was performed over an 85-min run at a flow rate of 0.4 mL/min using water containing 0.1% FA as mobile phase A and acetonitrile containing 0.1% FA as mobile phase B. Mass spectra were acquired in positive-ion mode over an m/z range of 100–1700, with MS/MS spectra acquired over an m/z range of 25–1000. The source temperature was 325 °C, with a drying-gas flow of 13 L/h and nebulizer pressure of 35 psi. Capillary and nozzle voltages were 4000 and 500 V, respectively. Collision energies were calculated as 3.1 × (m/z/100) + 1 for singly and doubly charged ions and 3.6 × (m/z/100) - 4.8 for ions with charge states ≥3. A reference mass of m/z 922.0098 was used for mass calibration.

### Proteomic data processing and quantitative analysis

Raw Agilent data files (.d) were converted to mzXML format using MSConvert and OpenMS. Protein identification and label-free quantification were performed using MaxQuant version 2.6.3 against the UniProt reference proteome for *Drosophila melanogaster* (taxonomy ID 7227), with trypsin specified as the digestion enzyme. Carbamidomethylation of cysteine was set as a fixed modification, whereas N-terminal acetylation and methionine oxidation were specified as variable modifications. The minimum LFQ ratio count was set to 1, and peptide-spectrum match and protein false-discovery rates (FDR) were set to 0.1. Protein groups identified only by site, corresponding to reverse sequences, or classified as potential contaminants were excluded, yielding 260 protein groups for downstream analysis. LFQ intensities were normalized during quantitative analysis and imported into MetaboAnalyst v6.0 for preprocessing and multivariate analysis. Features with all-zero or single non-zero values were removed, and remaining zeros were imputed as one-fifth of the minimum positive value for each feature, followed by log10 transformation. No additional normalization was performed. Pareto scaling was applied before principal component analysis (PCA) and unsupervised hierarchical clustering. Statistical comparisons and criteria for defining differentially abundant proteins (DEPs) are described in the Statistical analysis section. Functional enrichment analysis of DEPs was performed using the Database for Annotation, Visualization and Integrated Discovery (DAVID), including Gene Ontology (GO) biological process, cellular component, and molecular function categories, as well as KEGG and Reactome pathways.

### Metabolomics sample preparation

For metabolomic profiling, 38 *Drosophila* heads were collected per biological replicate from control, neuronal ERO1L knockdown (ERO1L-IR), and neuronal ERO1L overexpression (ERO1L-OE) groups. Five biological replicates were analyzed per condition. Samples were stored at -80 °C until extraction and thawed on ice for 5-10 min before processing. Metabolites were extracted using 200 µL methanol and 85 µL HPLC-grade water per sample. An extraction blank was included for quality control. Samples were thoroughly vortexed and sonicated for four 15-min cycles at 25 °C. Chloroform (200 µL) and HPLC-grade water (100 µL) were subsequently added, and samples were centrifuged twice at 15,000 × g for 15 min at 4 °C to achieve phase separation. The aqueous phase containing polar metabolites was collected and dried overnight at 40 °C using a centrifugal vacuum concentrator. Dried extracts were reconstituted in 300 µL of 50% acetonitrile in HPLC-grade water, filtered through 0.2-µm syringe filters, and transferred to LC–MS vials. A pooled quality-control (QC) sample was generated by combining 50 µL aliquots from each biological samples and serially diluted at 1:2, 1:4, 1:8, and 1:16.

### UHPLC-ESI-QTOF-MS/MS analysis

Metabolomic profiling was performed by ultra-high-performance liquid chromatography coupled to electrospray ionization quadrupole time-of-flight mass spectrometry (UHPLC-ESI-QTOF-MS/MS; Bruker, Germany). Chromatographic separation was performed using a Bruker Intensity Solo C18 column (2.1 × 100 mm, 2 µm) maintained at 40 °C, with the autosampler maintained at 4 °C. Mobile phase A consisted of water containing 0.1% FA, and mobile phase B consisted of acetonitrile containing 0.1% FA. The gradient consisted of 99% A from 0-2 min, 1% A from 2-20 min, 99% A from 20.1-28.3 min, and 99% A from 28.5-30 min. The flow rate was 0.25 mL/min, except from 20.1-28.3 min, when it was increased to 0.35 mL/min. The injection volume was 2 µL. Data were acquired separately in positive- and negative-ionization modes using a Bruker compact ESI-QTOF mass spectrometer with broadband collision-induced dissociation (bbCID), with the acquisition range of m/z 50-1300. External mass calibration was performed using a sodium formate solution containing 2 mM sodium hydroxide, 0.1% formic acid and 50% isopropanol, infused at 0.5 µL/min. The acquisition range was m/z 50-1300. Source conditions were optimized for high-resolution detection of metabolites in *Drosophila* head extracts. For positive-ionization acquisition, the cone voltage was set to 35 V and the capillary voltage to 4000 V. The source and desolvation temperatures were both maintained at 220 °C, with a desolvation-gas flow of 8 L/min. For negative-ionization acquisition, the cone voltage was set to 31 V and the capillary voltage to 4500 V, while the source and desolvation temperatures were maintained at 220 °C with a desolvation-gas flow of 8 L/min.

### Metabolomics data processing and metabolite annotation

Raw LC-MS data were processed using MS-DIAL for feature detection, alignment, quality-control (QC)-based signal correction, and metabolite annotation against the corresponding metabolite databases. Metabolite annotations were classified according to the level of assignment (LoA) used by the analytical facility: LoA 1, accurate-mass match to a database entry; LoA 2, accurate-mass and MS/MS match to in silico fragmentation patterns; LoA 3, MS/MS match to reference spectra from databases or the literature; LoA 4, retention-time and molecular-mass match to an authentic standard; and LoA 5, MS/MS match to an authentic standard. Features without an assigned LoA were designated as NA. Positive- and negative-ionization datasets were analyzed separately using MetaboAnalyst v6.0. No additional normalization or data transformation was applied, and feature intensities were Pareto scaled before multivariate analysis. Principal component analysis (PCA) was used to assess global metabolic variation. Duplicate and unannotated features were removed before pathway analysis. Annotated metabolites retained after feature selection were mapped to *Drosophila melanogaster* KEGG pathways using MetaboAnalyst v6.0. Metabolite set enrichment analysis and pathway topology analysis were performed to identify metabolic pathways associated with neuronal ERO1L modulation. Pathways were interpreted based on enrichment significance and pathway impact, and network analysis was used to visualize relationships among enriched pathways.

### Statistical analysis

Statistical analyses were performed using GraphPad Prism 11 (GraphPad Software, Boston, MA, USA). All tests were two-sided, and *P* < 0.05 was considered statistically significant. Data are presented as mean ± SEM or as box-and-whisker plots showing individual biological replicates and the minimum-to-maximum range, as indicated in the figure legends. Exact sample sizes, statistical tests, test statistics, and adjusted *P* values are reported in the corresponding figure legends. Biological replicates were independently generated experimental samples. For RT-qPCR and immunoblotting, biological replicates corresponded to independently prepared samples. For DHE staining and HP1 immunofluorescence, each independently dissected brain was considered one biological replicate. For HP1 quantification, measurements from 50 nuclei per brain were averaged to generate a single biological-replicate value; nuclei were treated as technical subsamples rather than independent observations. In negative-geotaxis assays, the mean climbing index across five consecutive trials was calculated for each vial, and each vial was treated as an independent experimental unit. For comparisons between two groups, unpaired two-tailed *t*-tests with Welch’s correction were used where appropriate. Comparisons among three groups were performed using one-way ANOVA followed by the relevant multiple-comparisons test; Welch’s ANOVA followed by Games-Howell testing was used when variances were unequal. For factorial experiments, two-way ANOVA was used to assess main effects and interactions, followed by the relevant post hoc multiple-comparisons test. The statistical test used for each experiment is specified in the corresponding figure legend. Proteomic LFQ data were compared using unpaired two-tailed *t*-tests with Welch’s correction. Differentially abundant proteins were defined using a fold-change threshold of ≥2 or ≤0.5 and nominal *P* < 0.1. For metabolomic data, partial least-squares discriminant analysis (PLS-DA) was performed separately for positive- and negative-ionization datasets. Model performance was evaluated by 5-fold cross-validation using one to five components, with Q² used to assess predictive performance and select the optimal number of components. Four components yielded the highest Q² in both ionization modes. Metabolite features contributing to group discrimination were ranked according to variable importance in projection (VIP), and features with VIP >1 were retained for downstream pathway analysis. Metabolomic pathway and functional-enrichment analyses were performed using the respective analytical platforms, and nominal enrichment P values were considered exploratory. No adjustment for multiple testing was applied to the proteomic or exploratory pathway analyses; interpretation therefore considered effect size, pathway-level convergence, and consistency across datasets. No data were excluded from statistical analyses except samples that failed predefined technical-quality criteria, including unsuccessful dissection, inadequate image quality, insufficient sample material, or assay-specific quality-control failure. Any exclusions and their reasons are reported in the relevant figure legends.

## Results and Discussion

### Neuronal ERO1L elevation reveals a pre-symptomatic stage before oxidative stress and locomotor decline

We previously found that *ERO1L* expression increases in aging *Drosophila* heads and that neuronal *ERO1L* elevation shortens lifespan under conditions of neuronal proteotoxic stress (Yeewa et al., 2024). To determine whether the age-associated increase in head *ERO1L* expression was preserved in the genetic background used for the present study, we quantified *ERO1L* transcript abundance in *y* flies, the background strain for the UAS-*ERO1L* stock. *ERO1L* expression remained broadly stable between 7 (D7) and 21 (D21) days post-eclosion but increased 2.19-fold by day 28 (D28) and continued to rise through day 42 (D42) (Fig. 1A). Thus, the increase in head *ERO1L* expression is not restricted to the previously examined *w^1118^* background but is reproduced in an independent genetic background.

**Figure 1.**
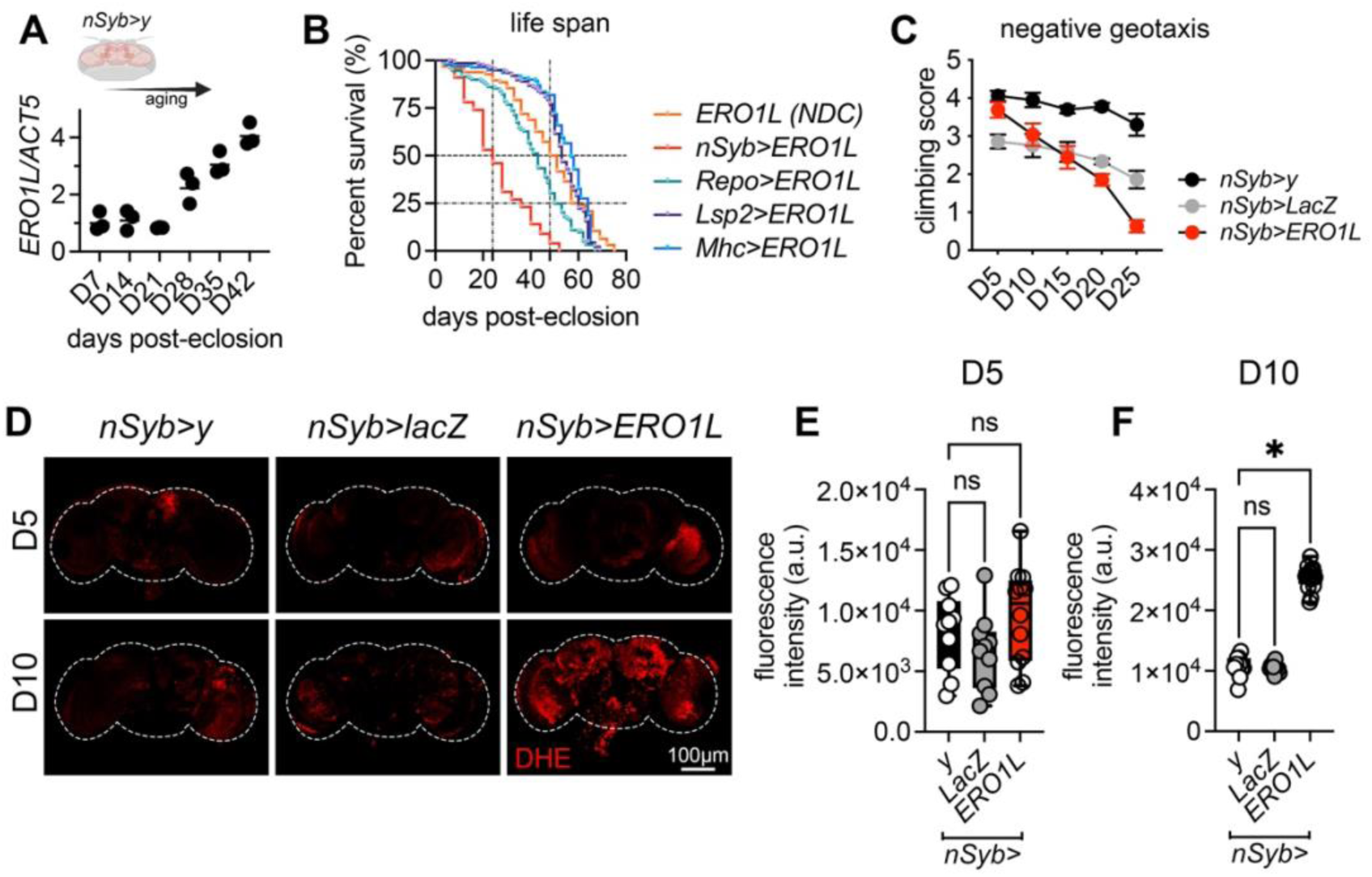
A presymptomatic window precedes neuronal ERO1L-induced oxidative stress and locomotor decline in *Drosophila*. **(A)** *ERO1L* mRNA abundance relative to *Act5C* in fly heads collected at the indicated days post-eclosion (control flies neuronal synaptobrevin (*nSyb)>y*). Endogenous head *ERO1L* expression increased with age. RNA was independently extracted from 50 heads per biological replicate; *n* = 3 biological replicates per time point. Each point represents one biological replicate. Scatter plot shows geometric mean. One-way ANOVA showed a significant effect of age on *ERO1L* expression [F(5,12)=36.08,P<0.0001]. Variance homogeneity was supported by the Brown–Forsythe test [F(5,12)=0.4342,P=0.8164]. **(B)** Kaplan–Meier survival curves of flies expressing *UAS-ERO1L* in neurons (*nSyb*>*ERO1L*), glia (*Repo*>*ERO1L*), muscle (Myosin heavy chain (*Mhc)*>*ERO1L*), or fat body (larval serum protein 2 (*Lsp2)*>*ERO1L*), compared with the corresponding non-driver control (ERO1L; NDC). Neuronal *ERO1L* expression markedly reduced lifespan, with a median survival of 24.0 days (95% CI, 20.0–28.0 days), compared with 49.5 days (95% CI, 45.0–54.0 days) in the NDC group. Glial *ERO1L* expression produced a substantially milder survival phenotype, with a median lifespan of 43.0 days (95% CI, 39.0–46.0 days). In contrast, *ERO1L* expression in muscle or fat body did not reduce survival, with median lifespans of 57.0 days (95% CI, 54.0–60.0 days) for *Mhc*>ERO1L and 53.0 days (95% CI, 52.0–56.0 days) for *Lsp2*>ERO1L flies. Cohort sizes were: *ERO1L*; NDC, *n* = 96; *nSyb*>*ERO1L*, *n* = 100; *Repo*>*ERO1L*, *n* = 122; *Mhc*>*ERO1L*, *n* = 125; and *Lsp2*>*ERO1L*, *n* = 131. **(C)** Negative-geotaxis performance of *nSyb*>y, *nSyb*>*LacZ*, and *nSyb*>*ERO1L* flies at the indicated ages. For each genotype and time point, 100 flies were assessed in 10 vials (10 flies per vial), with five consecutive climbing trials per vial; the mean climbing index across trials was used for statistical analysis.; data are presented as mean ± SEM. Neuronal *ERO1L* elevation caused an age-dependent decline in climbing performance. Two-way ANOVA revealed significant effects of age and genotype, as well as a significant age × genotype interaction (age: *F*(4,60) = 186.5, *P* < 0.000001; genotype: *F*(2,60) = 504.5, *P* < 0.000001; interaction: *F*(8,60) = 43.83, *P* < 0.000001). At D5, climbing performance did not differ significantly between *nSyb>ERO1L* and *nSyb>y* flies (Dunnett’s test; mean difference = 0.212, 95% CI, −0.033 to 0.456; adjusted *P* = 0.0977). From D10 onwards, *nSyb>ERO1L* flies showed significantly reduced climbing performance compared with *nSyb>y* control (all adjusted *P* < 0.000001), with the deficit becoming progressively greater with age. **(D)** Representative images of dihydroethidium (DHE) fluorescence in adult brains from *nSyb*>*y*, *nSyb*>*LacZ*, and *nSyb*>*ERO1L* flies at days 5 and 10 post-eclosion (D5 and D10, respectively). Dashed lines delineate the brain boundaries. No major increase in DHE fluorescence was observed in *nSyb*>*ERO1L* brains at day 5, whereas increased fluorescence was apparent at day 10. Scale bar, 100 µm. **(E–F)** Quantification of brain DHE fluorescence intensity at D5 (E) and D10 (F). Fluorescence intensity was quantified in independent brains from *nSyb*>*y*, *nSyb*>*LacZ*, and *nSyb*>*ERO1L* flies. Individual values are shown as box-and-whisker plots with minimum-to-maximum whiskers. At D5, DHE fluorescence did not differ significantly among genotypes (one-way ANOVA, F(2,29)=1.99, P=0.1554; Brown–Forsythe test, F(2,29)=0.835, P=0.4440; total n=32 brains). By day 10, DHE fluorescence differed significantly among groups (one-way ANOVA, F(2,35)=398, P<0.000001; Brown–Forsythe test, F(2,35)=1.62, P=0.2133; brains). Post-hoc multiple-comparisons analysis showed that DHE fluorescence was significantly increased in *nSyb*>*ERO1L* brains relative to both *nSyb*>*y* and *nSyb*>*LacZ* controls. ns, not significant; *P<0.05.

The timing of this increase is biologically relevant. *Drosophila* show progressive physiological decline across adult life, with deterioration of stress resistance, tissue function, metabolism, and locomotor capacity becoming increasingly evident during middle and late adulthood (Jones and Grotewiel, 2011; Piper and Partridge, 2018). Accordingly, D28 represents a transition into middle-aged adulthood in standard laboratory conditions, whereas D42 corresponds to a substantially aged stage. The progressive increase in head *ERO1L* abundance after D28 therefore suggests that *ERO1L* elevation accompanies later phases of brain aging.

The age-dependent increase in endogenous *ERO1L* expression provides a physiological context for the neuronal overexpression model. In control flies, head *ERO1L* expression increased progressively with age, reaching by D42 a magnitude similar to the approximately fourfold increase previously observed in *nSyb>ERO1L* flies (Yeewa et al., 2024). Thus, sustained pan-neuronal expression produces *ERO1L* levels within the range observed during late-life aging, supporting *nSyb>ERO1L* as a controlled approach to examine the consequences of sustained neuronal ERO1L activity at a magnitude relevant to late-life brain aging.

We next asked whether the lifespan-shortening effect of *ERO1L* elevation was specific to neurons. In agreement with our previous observations, pan-neuronal *ERO1L* expression driven by *nSyb*-GAL4 markedly reduced survival, with a median lifespan of 24 days compared with 49.5 days in the non-driver control (NDC) (Fig. 1B). In contrast, glial *ERO1L* expression driven by reversed polarity *Repo*-GAL4 produced a substantially milder phenotype, with a median lifespan of 43.5 days. This indicates that glial tissue can buffer sustained *ERO1L* elevation more effectively than neurons. *ERO1L* expression in muscle or fat body did not reduce survival. Median lifespan was 57 days in *Mhc*>*ERO1L* flies and 53 days in *Lsp2*>*ERO1L* flies, both exceeding the 49.5-day median of the NDC group (Fig. 1B). These findings indicate that the consequences of *ERO1L* elevation depend strongly on the cellular context, although differences in the level of *ERO1L* expression between tissues cannot be excluded as a contributing factor. The pronounced lifespan reduction observed with neuronal *ERO1L* elevation therefore identifies the nervous system as a particularly sensitive context for the detrimental effects of increased ERO1L activity.

Having established the neuron-selective toxicity of *ERO1L* elevation, we next determined when behavioural impairment first emerges. Negative-geotaxis performance declined progressively in *nSyb>ERO1L* flies with age (Fig. 1C). At day 5 post-eclosion (D5), however, their climbing scores remained comparable with those of both *nSyb>y* and *nSyb>LacZ* controls. By day 10 (D10), *nSyb>ERO1L* flies already showed reduced locomotor performance relative to control flies. Expression of *LacZ* also imposed a modest behavioural burden compared with the non-overexpressing *nSyb>y* control, consistent with the general cost of transgene expression; however, neuronal *ERO1L* elevation produced a more rapid and substantially larger decline in climbing performance.

Because ERO1 activity can promote oxidative protein folding and is linked to cellular redox imbalance and reactive oxygen species (ROS) production, we next assessed brain oxidative stress at the early behavioural time points. Dihydroethidium (DHE) fluorescence was measured in brains collected at D5, when locomotor performance remained preserved, and D10, when locomotor impairment was already evident. At D5, DHE fluorescence did not differ significantly between *ERO1L*-overexpressing brains and either control group (Fig. 1D–E). By D10, however, neuronal *ERO1L* elevation was associated with a marked increase in brain DHE fluorescence relative to the *nSyb>y* control (Fig. 1D, F).

Together, these data establish D5 as an operational pre-symptomatic stage of neuronal *ERO1L* elevation. At this time point, flies have not yet developed detectable locomotor impairment or major brain oxidative-stress accumulation, despite the later emergence of both phenotypes. Although several mechanisms have been proposed to mediate ERO1-associated cellular toxicity, including ER redox imbalance and altered ER–mitochondria functional coupling, the molecular consequences of chronically elevated neuronal ERO1L during this early functional window remain unknown. We therefore used D5 *nSyb>ERO1L* flies as a platform to define the initial molecular adaptations and vulnerabilities that precede overt neuronal dysfunction.

### Pre-symptomatic neuronal *ERO1L* elevation induces selective proteomic remodelling

We next asked whether sustained neuronal *ERO1L* elevation produces molecular changes before major brain oxidative stress and locomotor impairment. We performed label-free quantitative proteomics on D5 heads pan-neuronally expressing *UAS-ERO1L* (ERO1L-OE) flies and age-matched controls. The *y* control shares the genetic background of the *UAS-ERO1L* stock, minimizing potential effects of background variation (Fig. 2A, Supplementary Fig. 1A).

**Figure 2.**
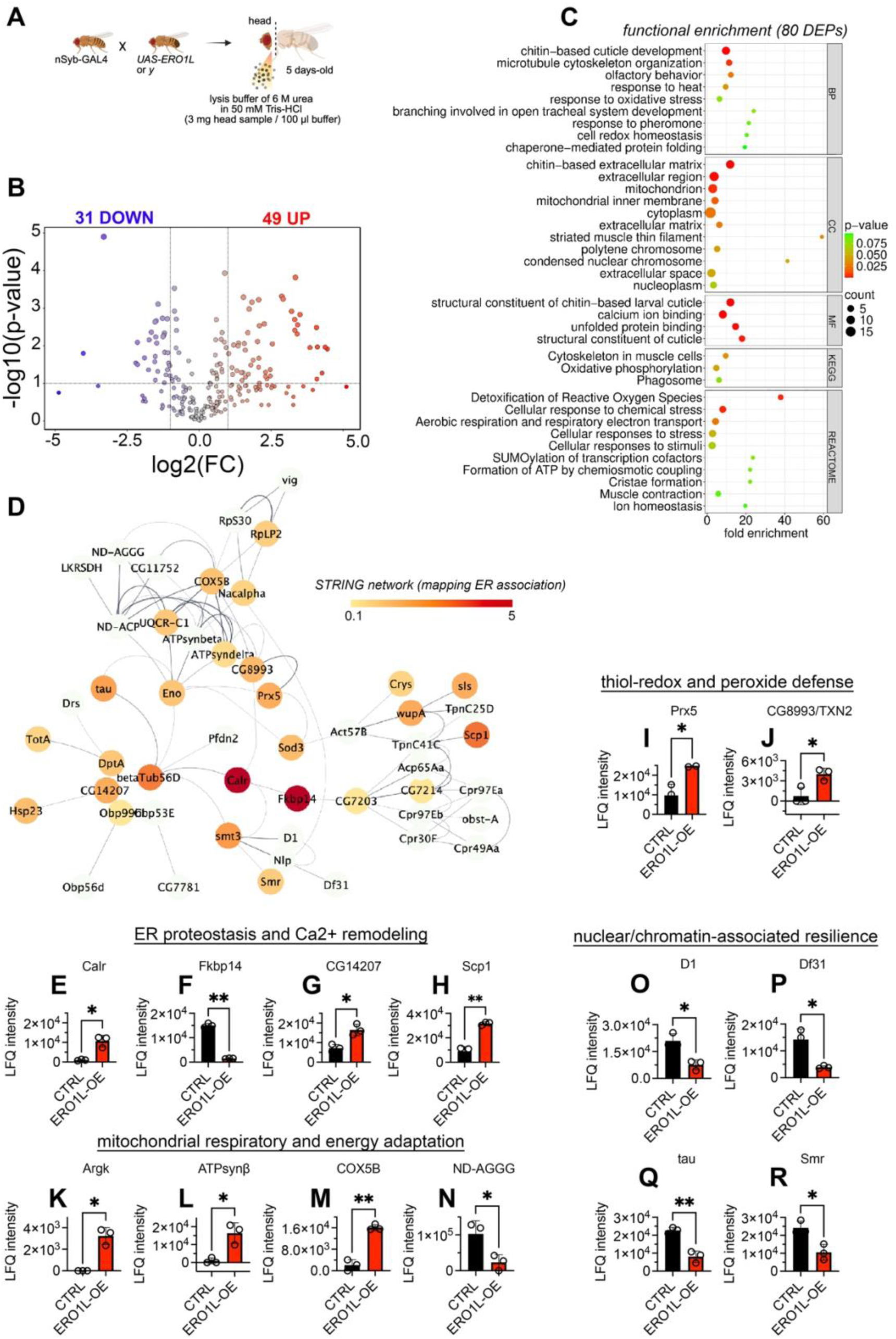
Presymptomatic neuronal *ERO1L* elevation induces coordinated ER–redox–bioenergetic adaptation and changes in chromatin-associated proteins. **(A)** Experimental workflow. *nSyb*-GAL4 flies were crossed with UAS-*ERO1L* to generate neuronal *ERO1L*-overexpressing flies (ERO1L-OE) or matched control (*nSyb>y*, CTRL). Heads were collected from 5-day-old adults and processed for label-free quantitative proteomic analysis. **(B)** Volcano plot showing differential protein abundance in ERO1L-OE relative to control heads. Each point represents one quantified protein. Differentially abundant proteins were defined as those showing a fold change ≥2 or ≤0.5 and a nominal *P* < 0.1. Red and blue points indicate increased and decreased protein abundance, respectively. A total of 31 proteins were decreased and 49 were increased. Dashed lines indicate the fold-change and *P*-value thresholds. **(C)** Functional enrichment analysis of the 80 differentially expressed proteins (DEPs). Enriched biological process (BP), cellular component (CC), molecular function (MF), Kyoto Encyclopedia of Genes and Genomes (KEGG), and Reactome terms. Dot size indicates the number of proteins associated with each term, and dot color indicates the nominal enrichment *P* value. **(D)** STRING functional-association network of DEPs, visualized in Cytoscape using the yFiles Radial Layout. Node color represents the ER-association mapping score, with lighter to darker red indicating lower to higher scores, respectively. Edges represent STRING functional associations. **(E–H)** Label-Free Quantification (LFQ) intensity profiles of proteins associated with ER proteostasis and calcium homeostasis: calreticulin (Calr), FK506-binding protein 14 (Fkbp14), heat shock protein B8/CG14207 (CG14207), and sarcoplasmic calcium-binding protein 1 (Scp1). **(I–J)** LFQ intensity profiles of proteins involved in thiol redox regulation and peroxide defense: peroxiredoxin 5 (Prx5) and thioredoxin 2 (CG8993/TXN2). **(K–N)** LFQ intensity profiles of proteins associated with mitochondrial respiration and energy metabolism: arginine kinase 1 (ArgK), ATP synthase subunit β (ATPsynβ), cytochrome c oxidase subunit 5B (COX5B), and NADH dehydrogenase accessory subunit ND-AGGG. **(O–R)** LFQ intensity profiles of proteins associated with nuclear/chromatin regulation and neuronal structural resilience: D1 chromosomal protein (D1), decondensation factor 31 (Df31), tau, and SMRTER corepressor (Smr). Circles represent individual biological replicates. Statistical comparisons of LFQ intensity were performed using unpaired two-tailed *t*-tests with Welch’s correction. \**P* < 0.05; \*\**P* < 0.01.

Principal component analysis (PCA) separated ERO1L and control samples, with PC1 accounting for 61.5% of total variance (Supplementary Fig. 1B). Unsupervised hierarchical clustering similarly grouped samples by genotype (Supplementary Fig. 1C), indicating a reproducible proteomic response to neuronal *ERO1L* elevation at D5. The complete LC–MS/MS dataset is provided as Electronic Supplementary Material 1.

Using a fold-change threshold of ≥2 or ≤0.5 and nominal P < 0.1, we identified 80 differentially abundant proteins, including 31 decreased (DOWN) and 49 increased (UP) proteins (Fig. 2B; Supplementary Table 3). Functional enrichment did not identify a single dominant pathway but instead revealed interconnected processes involving protein quality control, redox regulation, calcium handling, and energy metabolism. Biological-process terms included responses to heat, oxidative and chemical stress, chaperone-mediated protein folding, and redox homeostasis. Molecular-function analysis highlighted unfolded-protein and calcium-ion binding, while KEGG and Reactome analyses identified oxidative phosphorylation, ROS detoxification, cellular respiration, and ATP synthesis (Fig. 2C).

This pattern is consistent with the role of ERO1L as an ER oxidoreductase that supports oxidative protein folding while generating H₂O₂. The enrichment of protein-folding and redox-defence pathways suggests that sustained *ERO1L* elevation engages homeostatic mechanisms capable of buffering increased ER and redox demands rather than causing widespread proteome alteration.

To examine the organization of these responses, we generated a STRING functional-association network and visualized it in Cytoscape using the yFiles Radial Layout (Shannon et al., 2003), overlaying ER-association mapping scores (Fig. 2D). The network connected proteins involved in stress responses, redox regulation, mitochondrial metabolism, cytoskeletal organization, and nuclear functions. Notably, only calreticulin (Calr) and FK506-binding protein 14 (Fkbp14) showed strong ER-association scores. Thus, despite ERO1L being an ER-resident oxidoreductase, the early proteomic response extended beyond the ER to interconnected systems.

We next examined representative proteins from the major functional patterns identified by the enrichment and network analyses. These included ER proteostasis and calcium-associated proteins, thiol-redox and peroxide-defence proteins, mitochondrial respiratory and energy-associated proteins, and nuclear/chromatin-associated proteins (Fig. 2E-R).

Within the ER-associated module, Calr and CG14207 were increased, whereas Fkbp14 was reduced in ERO1L-overexpressing heads (Fig. 2E–G). Calreticulin is an ER lectin chaperone involved in glycoprotein quality control and ER luminal Ca²⁺ buffering (Schurch et al., 2022), Fkbp14 is an ER-associated foldase (Baumann et al., 2012), and CG14207 is a small heat-shock-protein family member associated with protein folding (Vos et al., 2016). The reciprocal regulation of these proteins suggests selective remodelling of ER proteostasis rather than uniform induction of folding factors. Sarcoplasmic calcium-binding protein 1 (Scp1), a calcium-binding protein, was also increased (Fig. 2H), further implicating calcium-associated processes in the early ERO1L response. Given the established role of ERO1 in regulating ER-to-mitochondria Ca²⁺ signalling, these changes support a model in which neuronal *ERO1L* elevation influences calcium homeostasis, in line with the known coupling between ER redox state and Ca²⁺ signalling.

The redox module showed a similarly coordinated response. Peroxiredoxin 5 (Prx5) and CG8993, the *Drosophila* ortholog of human thioredoxin 2 (TXN2), were increased (Fig. 2I–J). Prx5 participates in peroxide detoxification (Radyuk et al., 2010), whereas TXN2 contributes to thiol-redox regulation (Svensson and Larsson, 2007). Their increased abundance is consistent with the enrichment of ROS-detoxification and redox-homeostasis pathways and suggests that antioxidant capacity is engaged before overt oxidative stress. The increase in redox-defence proteins at D5 coincided with the absence of detectable ROS accumulation, consistent with an early antioxidant response that could help buffer the oxidative burden imposed by elevated *ERO1L*. This response may become insufficient with sustained *ERO1L* elevation, as increased brain ROS was evident by D10.

Mitochondrial and energy-associated proteins were also altered, consistent with the functional coupling between ER redox signalling, calcium homeostasis, and mitochondrial metabolism. Given the established connection between ERO1L and ER-mitochondria signalling (Anelli et al., 2012; Bassot et al., 2023), we generated a STRING network using mitochondrial association as the mapping criterion, which identified a broader mitochondrial-associated network (Supplementary Fig. 2B). Within this network, Cytochrome c oxidase subunit Vb (COX5B), ATP synthase beta subunit (ATPsynβ), and Arginine kinase (ArgK) were increased, whereas NADH dehydrogenase (ubiquinone) 1 alpha subcomplex subunit GGG (ND-AGGG) was reduced (Fig. 2K–N). These proteins participate in oxidative phosphorylation, ATP synthesis, and cellular ATP buffering, respectively (Ajoolabady et al., 2022; Deng et al., 2018; Justs et al., 2026), suggesting selective remodelling of mitochondrial energy metabolism. Other mitochondrial-associated proteins were unchanged, including Adult cuticle protein 65Aa (Acp65Aa), Actin 57B (Act57B), CG11752, and NADH-ubiquinone oxidoreductase acyl-carrier protein; also called mitochondrial acyl carrier protein 1 (ND-ACP), whereas Odorant-binding protein 56d (Obp56d) was reduced (Supplementary Fig. 2C). Mitochondrial-associated singleton proteins were also heterogeneous, including increased Ferritin 2 light chain homologue (Fer2LCH), a protein involved in iron storage and oxidative-stress regulation (Tang and Zhou, 2013), and decreased Haspin (Hasp), together with changes in Obstructor-E (Obst-E), Salivary gland secretion 3 (Sgs3), Slippery little kid (Slik), and Vasa intronic gene (vig) (Supplementary Fig. 2D). Thus, the proteomic response included a selective set of changes in proteins linked to mitochondrial respiration, ATP production, and metabolic homeostasis.

An unexpected feature of the D5 proteome was the reduction of proteins associated with nuclear and chromatin-related functions. D1 chromosomal protein (D1), Decondensation factor 31 (Df31), Tau, and SMRT-related ecdysone receptor-interacting protein (Smr) were reduced in ERO1L-OE heads (Fig. 2O–R). Because these proteins participate in chromatin organization and nuclear or neuronal structural functions (Bolkan and Kretzschmar, 2014; Heck et al., 2012; Schubert et al., 2012; Smith and Weiler, 2010), their reduction suggests that nuclear homeostasis may become vulnerable even as other cellular systems engage compensatory responses. Overall, the D5 proteome shows that neuronal *ERO1L* elevation does not cause widespread proteome disruption but instead triggers a selective response involving ER proteostasis, redox defence, calcium regulation and mitochondrial energy metabolism. Changes in proteins linked to energy production and metabolism further suggest that the response extends beyond proteostasis and redox control to affect neuronal metabolism. These adaptations occur while locomotor function and bulk brain ROS remain largely preserved. At the same time, the selective reduction of nuclear- and chromatin-associated proteins points to an early vulnerability within this otherwise adaptive response. Thus, D5 represents a pre-symptomatic state in which neuronal homeostasis is actively maintained, while changes in nuclear- and chromatin-associated proteins point to an additional layer of the cellular response.

### Neuronal *ERO1L* elevation is accompanied by remodelling of central-carbon, nitrogen, and purine metabolism

Next, we asked whether the remodelling of metabolic proteins observed at the pre-symptomatic stage was accompanied by corresponding changes at the metabolite level. To place the ERO1L phenotype in the context of altered ERO1L abundance, we also included neuronal *ERO1L* knockdown (ERO1-IR) in the analysis. Genetic reduction of *ERO1L* has previously been shown to extend lifespan in *Drosophila (Yeewa et al., 2024)*, providing a contrasting perturbation to *ERO1L* elevation. Comparing ERO1L-OE and ERO1-IR with the same control (CTRL) allowed us to determine whether metabolic changes were specific to increased *ERO1L* or reflected a broader response to altered *ERO1L* abundance.

Untargeted metabolomic profiling was performed in positive- and negative-ion acquisition modes (Fig. 3A, Supplementary Fig. 3A). Partial least-squares discriminant analysis (PLS-DA) clearly separated ERO1L-OE from CTRL samples in both modes, indicating a reproducible metabolic shift associated with elevated neuronal ERO1L (Supplementary Fig. 3B–E). In contrast, ERO1L-IR samples remained closely associated with CTRL in positive-ion mode and showed partial overlap with CTRL in negative-ion mode. Thus, the major metabolic shift at D5 was associated with *ERO1L* elevation rather than reduction, indicating that increased *ERO1L* has a greater impact on the head metabolome than its suppression under these experimental conditions. Complete feature tables for both acquisition modes are provided as Electronic Supplementary Material 2 and 3.

**Figure 3.**
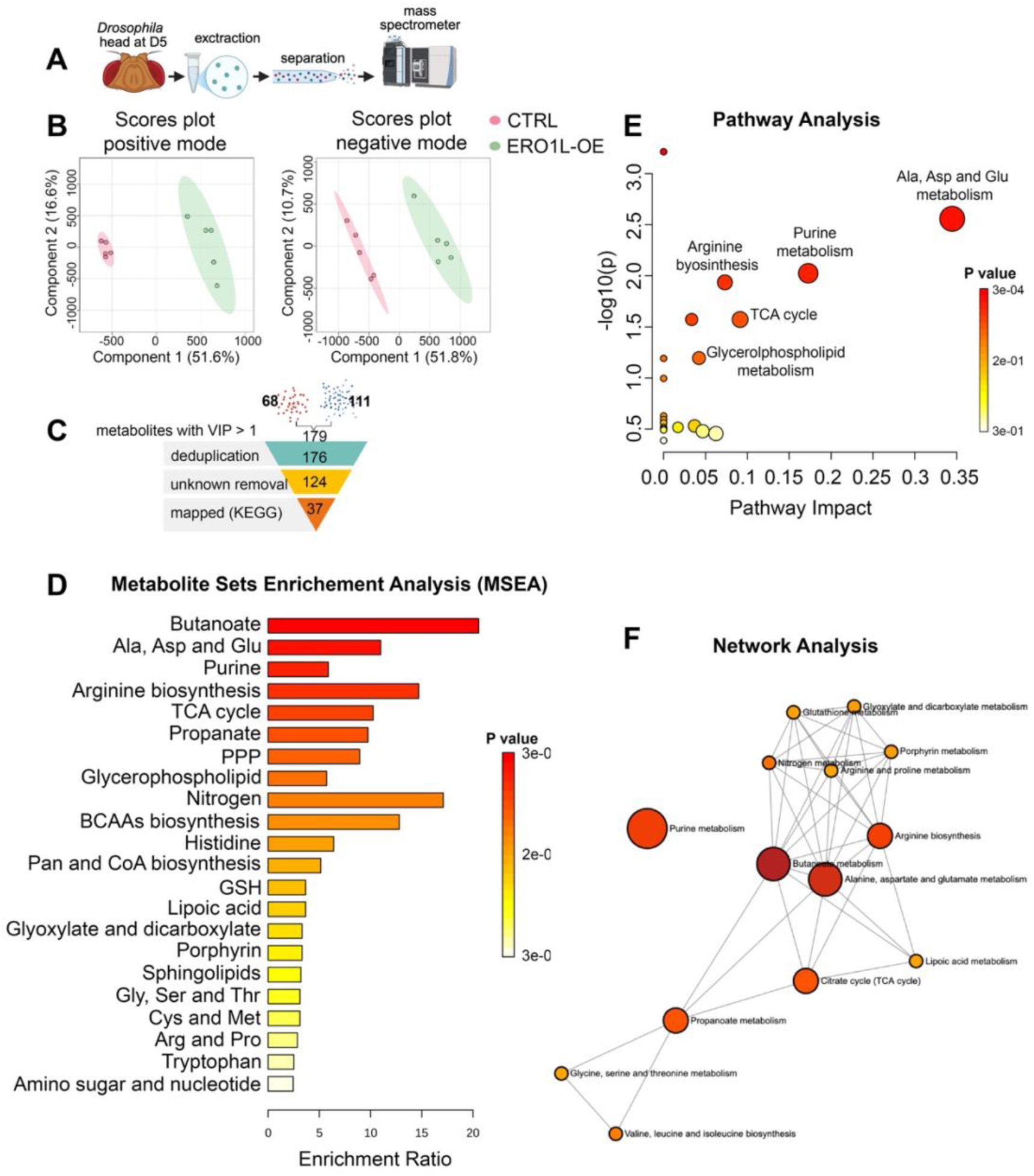
Pre-symptomatic neuronal *ERO1L* elevation is associated with remodeling of central-carbon, amino-acid, nitrogen, and purine metabolism. **(A)** Experimental workflow for untargeted metabolomic profiling of 5-day-old control (CTRL) and neuronal *ERO1L*-overexpressing (ERO1L-OE) fly heads, under pan-neuronal nSyb-GAL driver. Metabolites were extracted using methanol/chloroform and analyzed in positive- and negative-ion modes. **(B)** Partial least-squares discriminant analysis score plots of CTRL and ERO1L-OE head metabolomes acquired in positive-ion mode (left) and negative-ion mode (right). ERO1L-OE and CTRL samples segregated primarily along component 1, which explained 51.6% and 51.8% of the variance in positive- and negative-ion modes, respectively. Component 2 explained 16.6% and 10.7% of the variance, respectively. Each point represents one biological replicate; shaded ellipses indicate group distributions. **(C)** Selection and annotation workflow for discriminatory metabolite features. A total of 179 features with variable-importance-in-projection (VIP) scores >1 were identified, including 68 and 111 features preferentially associated with the two experimental groups. After deduplication and removal of unknown features, 37 metabolites were mapped to KEGG pathways. **(D)** Metabolite-set enrichment analysis (MSEA) of discriminatory metabolites. Pathways are ranked by enrichment ratio. Bar length represents enrichment ratio and color denotes enrichment P value. **(E)** Pathway-impact analysis of differential metabolites. The x-axis indicates pathway impact and the y-axis indicates -log10(P); color denotes pathway-enrichment P value. **(F)** Network analysis of enriched metabolite sets. Nodes represent metabolite sets, with node size proportional to enrichment ratio and node color indicating enrichment P value. Edges indicate relationships among metabolite sets.

Variable-importance-in-projection (VIP) analysis identified the features contributing most strongly to the separation between ERO1L-OE and CTRL samples in each acquisition mode (Supplementary Tables 4 and 5). After filtering for VIP scores >1 and removing duplicate and unannotated features, 37 metabolites were mapped to KEGG pathways (Fig. 3C; Supplementary Table 6). Enrichment analysis identified interconnected pathways involving central-carbon, amino-acid, nitrogen, purine, and lipid metabolism, rather than a single dominant metabolic pathway (Fig. 3D).

Pathway-topology analysis prioritized alanine/aspartate/glutamate metabolism, followed by purine metabolism, arginine biosynthesis, the citrate cycle, and glycerophospholipid metabolism (Fig. 3E). Network analysis further showed that these pathways were highly interconnected, linking amino-acid and nitrogen metabolism with central-carbon metabolism, mitochondrial energy pathways, nucleotide metabolism, and membrane-lipid metabolism (Fig. 3F). This organization is particularly relevant to the proteomic findings. ERO1L-OE heads showed increased COX5B, ATPsynβ, and ArgK and reduced ND-AGGG, indicating selective remodelling of respiratory-chain, ATP-synthesis, and ATP-buffering proteins (Fig. 2K–N). The enrichment of citrate-cycle, amino-acid, arginine, and nitrogen pathways therefore provides a biochemical counterpart to the mitochondrial and energy-associated proteomic response.

The connection between these datasets is also evident at the level of redox metabolism. The pentose-phosphate pathway was enriched in ERO1L-OE heads, providing a potential link to cellular reducing capacity through NADPH production. This finding accompanies increased Prx5 and CG8993/TXN2 in the proteome, consistent with coordinated engagement of metabolic and antioxidant systems. Purine metabolism provides a further connection to energy homeostasis, as purine nucleotides participate directly in cellular energy metabolism (Grychowski et al., 2026). Additional enrichment of glycerophospholipid, branched-chain amino-acid, butanoate, and propanoate metabolism indicates that the response extends beyond mitochondrial respiration to broader remodelling of carbon, amino-acid, and lipid metabolism.

Together, the pathway-level analyses place the metabolic response within the broader cellular consequences of sustained *ERO1L* elevation.

### Selective changes in amino-acid and purine metabolites accompany neuronal *ERO1L* elevation

To resolve pathway-level signatures into individual metabolites, we examined alanine/aspartate/glutamate and purine metabolism, two pathways highlighted by metabolite-set enrichment and pathway-topology analyses (Fig. 3D–F). ERO1L-OE heads displayed a selective amino-acid/TCA-linked metabolic signature. α-Ketoglutarate (α-KG) and 2-ketobutyrate (2-KB) were increased relative to both CTRL and ERO1L-IR heads (Fig. 4A,C,E; Supplementary Table 6), whereas L-glutamate was elevated in both ERO1L-modulated groups relative to CTRL, without differing between ERO1L-OE and ERO1L-IR heads (Fig. 4A,D; Supplementary Table 6). Thus, while glutamate homeostasis responded to altered neuronal *ERO1L* abundance irrespective of direction, elevated *ERO1L* was specifically associated with increased α-KG and 2-KB. Because α-KG is a citrate-cycle intermediate and acceptor in transamination reactions (Xiao et al., 2016), and 2-KB is an amino-acid-derived α-ketoacid (Bui et al., 2019), these changes are consistent with altered integration of amino-acid carbon metabolism, nitrogen handling, and mitochondrial metabolism.

**Figure 4.**
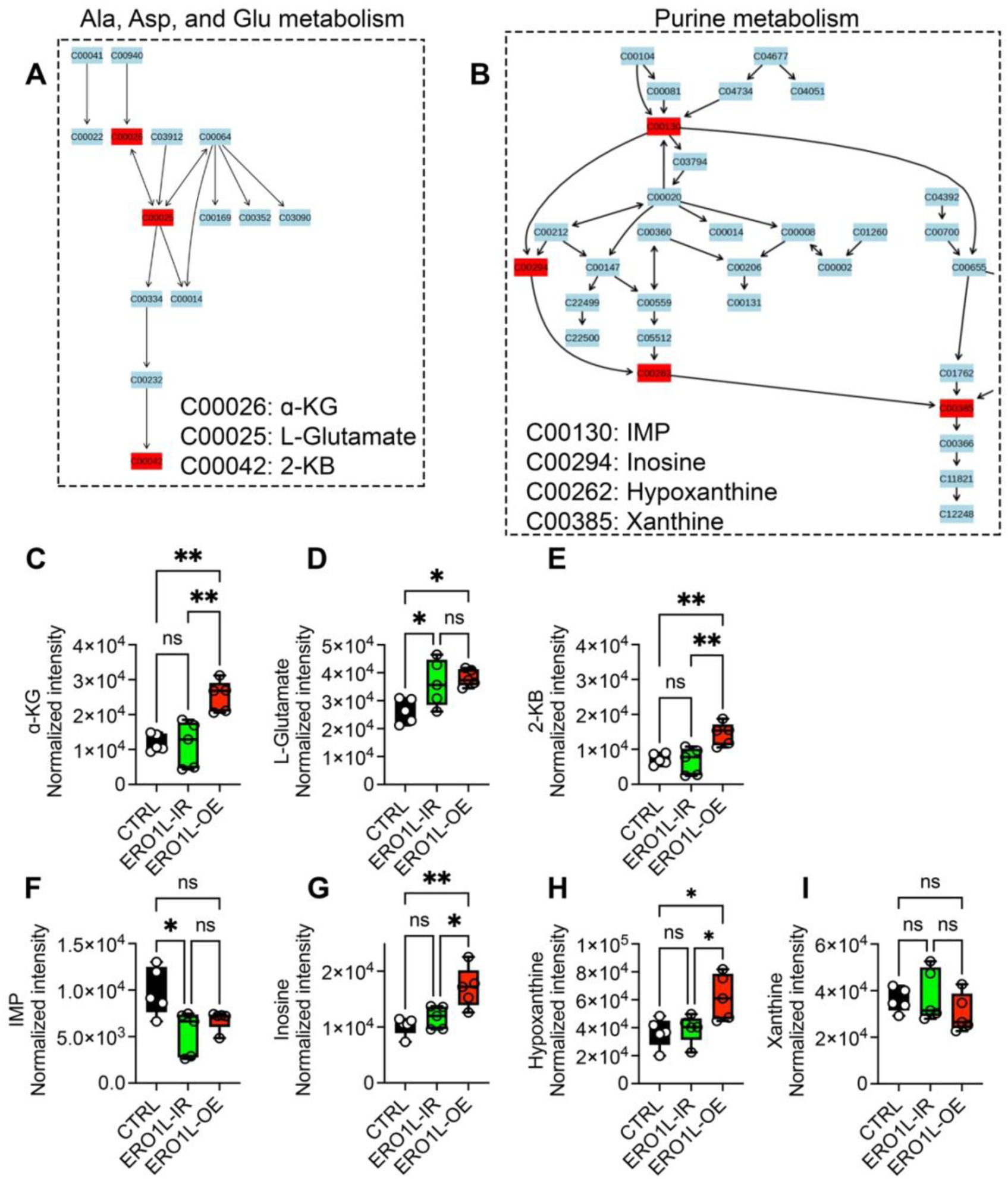
Neuronal *ERO1L* elevation is associated with selective remodelling of amino-acid and purine metabolism at the pre-symptomatic stage. **(A)** KEGG pathway map of alanine, aspartate, and glutamate metabolism. Differential metabolites identified in the analysis are highlighted in red, including alpha-ketoglutarate (α-KG; C00026), L-glutamate (C00025), and 2-ketobutyrate (2-KB; C00042). **(B)** KEGG pathway map of purine metabolism. Differential metabolites highlighted in red include inosine monophosphate (IMP; C00130), inosine (C00294), hypoxanthine (C00262), and xanthine (C00385). **(C–E)** Relative peak of α-KG; C00026 **(C)**, L-glutamate **(D)**, and 2-KB **(E)** in control (CTRL), neuronal *ERO1L* RNAi (ERO1L-IR), and neuronal *ERO1L*-overexpressing (ERO1L-OE) heads. α-KG and 2-KB were increased in ERO1L-OE relative to CTRL and ERO1L-IR. L-glutamate was increased in both ERO1L-IR and ERO1L-OE relative to CTRL, with no significant difference between ERO1L-IR and ERO1L-OE. **(F–I)** Normalized intensities of IMP **(F)**, inosine **(G)**, hypoxanthine **(H)**, and xanthine **(I)**. IMP was reduced in ERO1L-IR relative to CTRL, whereas inosine and hypoxanthine were increased in ERO1L-OE relative to both CTRL and ERO1L-IR. Xanthine abundance was not significantly altered among groups. Box-and-whisker plots show minimum-to-maximum values. Statistical comparisons were performed using one-way ANOVA followed by Tukey’s multiple-comparisons test. ns, not significant; *P<0.05; **P<0.01.

Purine metabolism showed a similarly selective response. Inosine and hypoxanthine were increased in ERO1L-OE heads relative to both CTRL and ERO1L-IR heads (Fig. 4B,G,H; Supplementary Table 6), whereas inosine monophosphate (IMP) was reduced only in ERO1-IR heads and xanthine was unchanged across groups (Fig. 4B,F,I; Supplementary Table 6). The absence of a uniform response across purine intermediates indicates remodelling of specific purine-metabolic nodes rather than generalized activation or depletion of the pathway.

The metabolomic profile complements the D5 proteomic signature of mitochondrial and energy-associated remodelling, including increased COX5B, ATPsynβ and ArgK and reduced ND-AGGG. Together, these changes indicate that neuronal ERO1L elevation is accompanied by early metabolic reorganization involving mitochondrial energy handling, amino-acid carbon metabolism and purine metabolism. This broad response is consistent with the capacity of neurons to adjust interconnected metabolic pathways while locomotor function remains preserved and brain DHE-detectable oxidative signal is not significantly increased.

The metabolic changes also raise a potential link between this adaptive response and the nuclear phenotype identified in the D5 proteome. α-KG is an obligate co-substrate for several dioxygenases involved in histone and DNA demethylation (Xu et al., 2011), whereas purine metabolism contributes to nucleotide availability for nucleic-acid synthesis, repair and transcription (Tran et al., 2024). The altered α-KG and purine profiles therefore provide a plausible metabolic context for the concurrent reduction of the nuclear- and chromatin-associated proteins D1, Df31, Smr and vig.

### Pre-symptomatic neuronal *ERO1L* elevation is associated with early chromatin- and RNA-homeostasis vulnerability

These observations prompted us to examine whether the early metabolic remodelling was accompanied by changes in nuclear homeostasis. We therefore assessed markers of heterochromatin structure and RNA-dependent transposable-element (TE) regulation: Heterochromatin Protein 1 (HP1); histone H3 lysine 9 di- and trimethylation (H3K9me2 and H3K9me3), repressive marks associated with heterochromatin; and *Drosophila* Fragile X Mental Retardation 1 (dFmr1) and P-element induced wimpy testis (Piwi), which participate in RNA- and small-RNA-mediated TE regulation.

Immunoblot analysis showed reduced HP1 and dFmr1 abundance in ERO1L-OE (*nSyb>ERO1L*) heads relative to *nSyb>LacZ* controls (Fig. 5A–C; Supplementary Fig. 4). Piwi abundance was higher than in the *y* control but did not differ significantly between *ERO1L-OE* and *LacZ* heads (Fig. 5A,D; Supplementary Fig. 4). We next examined H3K9 methylation to determine whether the reduction in HP1 was accompanied by alteration of a major heterochromatin-associated mark. H3K9me2 was reduced in *ERO1L-OE* heads, whereas H3K9me3 was unchanged (Fig. 5E–G; Supplementary Fig. 4). Thus, neuronal *ERO1L* elevation was associated with a selective reduction in HP1, dFmr1, and H3K9me2, rather than a uniform decrease in chromatin- or TE-regulatory factors.

**Figure 5.**
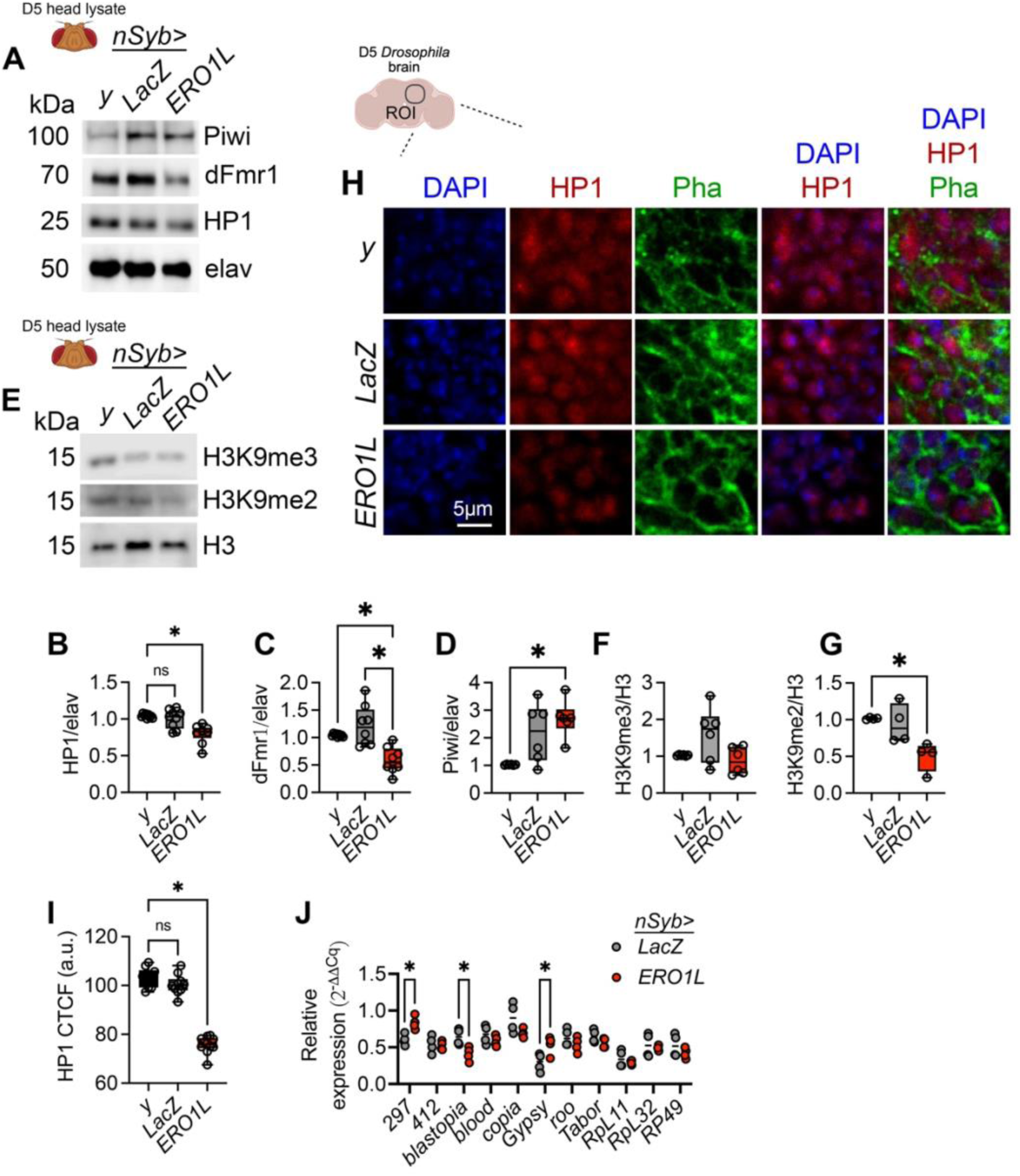
Neuronal ERO1L elevation is associated with early nuclear changes and selective transposable-element dysregulation. **(A)** Representative immunoblots of P-element-induced wimpy testis (Piwi), *Drosophila* fragile X mental retardation protein 1 (dFmr1), and heterochromatin protein 1a (HP1a) in head lysates from 5-day-old (D5) *y*, *nSyb*>*LacZ*, and *nSyb*>*ERO1L* flies. Embryonic lethal abnormal vision (Elav) was used as the normalization control. **(B–D)** Densitometric quantification of HP1 (B), dFmr1 (C), and Piwi (D), normalized to Elav. **(E)** Representative immunoblots of histone H3 lysine 9 trimethylation (H3K9me3) and histone H3 lysine 9 dimethylation (H3K9me2) in head lysates from D5 *y*, *nSyb*>LacZ, and *nSyb*>ERO1L flies. Total histone H3 was used as the normalization control. **(F–G)** Quantification of H3K9me3 (F) and H3K9me2 (G), normalized to total H3. **(H)** Representative HP1 immunofluorescence images of adult brains. DAPI, blue; HP1, red; phalloidin (Pha), green. Scale bar, 5 µm. A cartoon of the *Drosophila* brain indicates the central brain region used as the region of interest (ROI) for HP1 quantification. **(I)** Quantification of HP1 corrected total cell fluorescence (CTCF). **(J)** Relative abundance of the indicated TE transcripts in 5-day-old *nSyb>LacZ* and *nSyb>ERO1L* heads, measured by RT-qPCR and normalized to *GAL4*. RNA was extracted from 50 heads per biological replicate; *n* = 4 independent biological replicates per genotype. Data are shown as box-and-whisker plots with individual biological replicates and minimum-to-maximum range. For panels B–D and F–G, data were analyzed by Welch’s one-way ANOVA with heteroscedasticity-robust multiple-comparisons testing. HP1a was reduced in *nSyb>ERO1L* heads relative to *y* controls (*P* = 0.0010), dFmr1 was reduced relative to both *y* (*P* = 0.0023) and *nSyb>LacZ* (*P* = 0.0043) controls, and H3K9me2 was reduced relative to *y* controls (*P* = 0.0329); H3K9me3 was unchanged. Panel I was analyzed by one-way ANOVA with Šidák’s multiple-comparisons test; HP1 CTCF was reduced in *nSyb>ERO1L* brains relative to controls (*P* < 0.0001). Panel J was analyzed by two-way ANOVA with genotype and TE as factors, followed by Šidák-adjusted comparisons; the genotype-by-TE interaction was significant (*P* = 0.0003), with increased 297 (*P* = 0.0387) and gypsy (*P* = 0.0292) and reduced blastopia (*P* = 0.0321) in *nSyb>ERO1L* heads. ns, not significant; \**P* < 0.05.

To determine whether reduced HP1 abundance was also evident within the brain, we performed HP1 immunofluorescence. HP1 signal was significantly reduced in ERO1L-OE brains relative to control (Fig. 5H,I), confirming that the change detected in head extracts was also present in brain tissue.

We next assessed TE transcript abundance. Among the elements examined, *297* and *gypsy* transcripts were increased in *ERO1L-OE* heads, whereas *blastopia* was reduced and the remaining TEs were unchanged (Fig. 5J). Thus, sustained neuronal *ERO1L* elevation was associated with element-specific TE-expression changes, rather than broad activation of the examined TE panel. The increased expression of the LTR retrotransposon *gypsy* is notable because *gypsy* expression and activity increase in aging *Drosophila* brains, and aberrant *gypsy* activity has been associated with neurodegenerative phenotypes in fly models (Rigal et al., 2022; Wood et al., 2016).

Together, these findings identify chromatin- and RNA-homeostasis disruption as an early feature of the neuronal ERO1L response. The coordinated reduction in HP1, dFmr1 and H3K9me2, together with element-specific TE dysregulation, indicates that sustained *ERO1L* elevation is associated with selective weakening of mechanisms that support heterochromatin integrity and TE regulation. Given the established links between heterochromatin dysregulation, TE activation, brain aging and neurodegeneration, this nuclear signature provides a plausible molecular route through which chronic *ERO1L* elevation may contribute to later neuronal vulnerability.

## Conclusion

Sustained neuronal *ERO1L* elevation imposes a progressive, neuron-selective burden that is initially met by extensive molecular adaptation. Although *ERO1L* expression increased in aging fly heads and pan-neuronal *ERO1L* overexpression markedly shortened lifespan, D5 *nSyb>ERO1L* retained normal locomotor performance and showed no major increase in brain DHE fluorescence. This early preservation did not reflect molecular quiescence. Rather, at a level of neuronal *ERO1L* elevation broadly comparable to that observed in late-life heads, D5 *nSyb>ERO1L* flies exhibited coordinated coordinated remodelling of ER proteostasis, redox defence, calcium-associated processes, mitochondrial energy metabolism, and central-carbon, amino-acid, nitrogen, and purine metabolism.

The convergence of these changes suggests that neurons enter a compensated state in which metabolic and redox networks are reorganized to accommodate chronic ERO1L-associated stress. Increased folding- and antioxidant-associated proteins altered respiratory and ATP-buffering components, and the α-KG/2-KB signature collectively point to substantial energetic and biochemical investment in maintaining homeostasis. The emergence of increased brain ROS and impaired locomotion by D10 indicates that this early adaptive state is finite, and that sustained *ERO1L* elevation may progressively exceed the capacity of neuronal redox and metabolic buffering systems.

A notable finding was the parallel emergence of an unexpected nuclear phenotype. Reduced D1, Df31, Smr, and vig abundance in the D5 proteome was accompanied by lower HP1, dFmr1, and H3K9me2, together with selective dysregulation of TE transcripts, including *297* and *gypsy*. These observations identify chromatin- and RNA-homeostasis vulnerability as an early component of the neuronal response to *ERO1L* elevation. This connection is particularly compelling in view of the established association between heterochromatin erosion, TE dysregulation and neurodegeneration.

Our data therefore support a model in which chronic neuronal *ERO1L* elevation generates two concurrent states: a metabolically demanding adaptive programme that temporarily preserves neuronal function, and a developing chromatin vulnerability that may constrain long-term resilience. Whether chromatin remodelling is driven directly by altered ER redox signalling, emerges secondarily from metabolic and energetic pressure, or contributes causally to later dysfunction remains unresolved. Nevertheless, the convergence of α-KG and purine-metabolite changes with nuclear-protein loss and heterochromatin-associated alterations reveals a previously unrecognized intersection between ERO1L activity and chromatin biology.

This framework raises the possibility that chromatin-associated changes could serve as early indicators of ER-redox-driven brain aging. Future studies should define the extent to which chromatin instability is directly dependent on sustained ERO1L activity and dissect its bidirectional relationship with metabolic stress. In particular, it will be important to determine whether preserving chromatin stability reduces the metabolic burden imposed by *ERO1L* elevation, or whether limiting ERO1L-associated ER-redox and metabolic stress restores chromatin-associated homeostasis. Testing these relationships across proteotoxic, mitochondrial, and NDs models will establish the generality of this signature and may reveal strategies to preserve neuronal resilience.

## Data availability

All data supporting the findings of this study are provided in the main article and its Supplementary Information, including uncropped and unedited images of all blots and gels. Numerical source data underlying the graphs are available from Figshare (Lo Piccolo, 2026). Raw proteomic data generated in this study have been deposited in the JPOST Proteomics repository under accession numbers JPST004885. The complete proteomic and metabolomic datasets generated and analysed in this study are provided as Electronic Supplementary Material. Additional data supporting the findings of this study are available from the corresponding author upon reasonable request.

## Code availability

No custom code or scripts were generated for this study. Data analysis was performed using commercially available software and standard analysis tools as described in the Methods.

## Ethics statement

This study used *Drosophila melanogaster* and did not involve vertebrate animals or human participants. Experiments were conducted in accordance with the institutional policies of Chiang Mai University governing the use of invertebrate models.

## Materials availability

*Drosophila* stocks used in this study are available from the Bloomington Drosophila Stock Center or from the corresponding author upon reasonable request. All antibodies used in this study are commercially available, and primer sequences are provided in the Methods. Additional biological materials used in this study are available from the corresponding author upon reasonable request, subject to institutional and material-transfer requirements.

## AUTHOR CONTRIBUTIONS

Conceptualization: L.LP. and S.J.; Data curation: L.LP. and S.J.; Formal Analysis: L.LP., R.Y., M.S., J.P., P.S.; Funding acquisition; L.LP.; Investigation: L.LP., R.Y., P.P., C.P., M.S., J.K., W.J., P.S.; Methodology: L.LP., J.P.,P.S.; Project administration: L.LP.; Resources; L.LP and S.J; Software: L.LP., J.P, P.S.; Supervision; L.LP; Validation: L.LP. and S.J.; Visualization: L.LP; Writing – original draft: L.LP.; Writing – review & editing: All authors have reviewed and approved the final version of this manuscript.

## ACKNOWLEDGMENTS

This work was supported by the Fundamental Fund (FF) from the Science, Research and Innovation Promotion Fund (TSRI), Chiang Mai University, FY2568 under Contract No. 207569. This study was partially supported by the FF, FY2568 Contract No. 207573, the Faculty of Medicine, Chiang Mai University grant no FACMED 139/2567, Health Systems Research Institute (HSRI) under grants 66-124 and 68-060, the Mid-Career Research Grant (N42A670768) of the National Research Council of Thailand. We thank Mr. Papon Hitmool, Mr. Sakorn Phongchankhiao, and Mr. Tawan Munwanna (Department of Pharmacology, Faculty of Medicine, Chiang Mai University) for their technical and administrative assistance. We thank Bloomington Stock Center for providing fly strains, the Science and Educational Company Limited (SCIED) for technical support in confocal scanning electron microscopy.

