## supplementary data for "Pre-symptomatic proteomic and metabolomic profiling identifies compensated ER-redox-metabolic adaptation and early nuclear vulnerability in neuronal ERO1L toxicity"

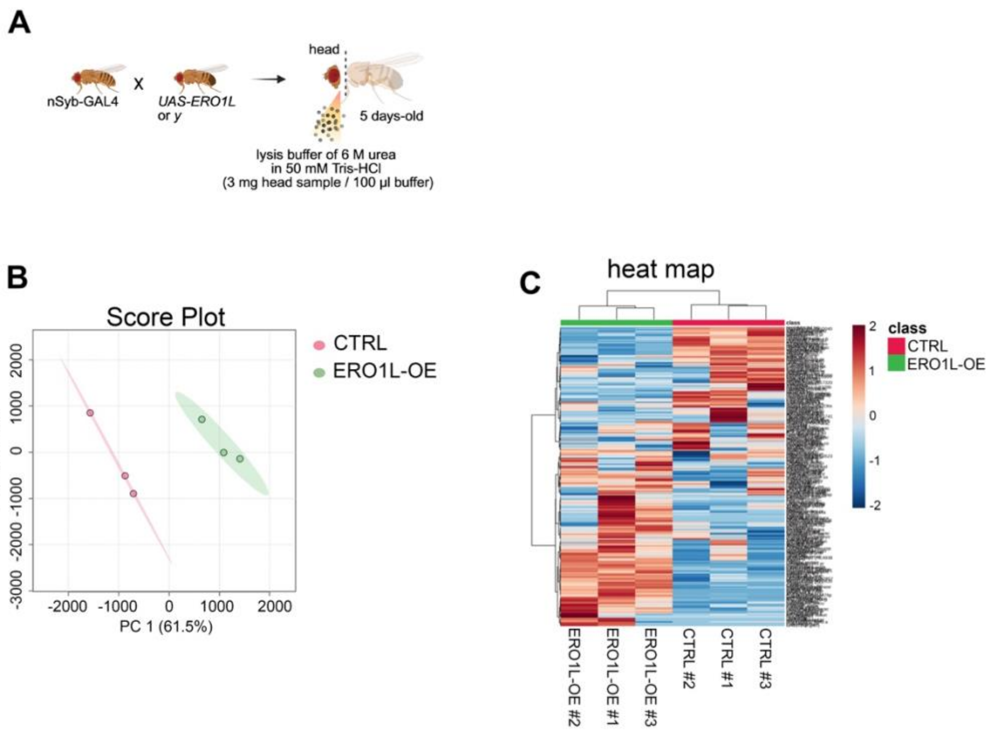


**Supplementary Figure 1 Experimental workflow and global proteomic separation of pre-symptomatic neuronal *ERO1L*-erexpressing heads.**
**(A)** Experimental workflow for label-free quantitative proteomics. nSyb-GAL4 flies were crossed to *UAS-ERO1L* to generate neuronal *ERO1L*-overexpressing flies (ERO1L-OE) or to the *y* control line (CTRL). Heads were collected from 5-day-old adults, corresponding to the pre-symptomatic stage, and lysed in 6 M urea/50 mM Tris-HCl buffer. Head lysates were prepared at 3 mg tissue per 100 µL lysis buffer and processed for proteomic analysis. **(B)** Principal-component analysis score plot of normalized label-free quantitative proteomic profiles from CTRL and ERO1L-OE head samples. CTRL and ERO1L-OE biological replicates segregated along principal component 1 (PC1, 61.5% of variance) and principal component 2 (PC2, 19.4% of variance), indicating distinct global proteomic profiles at the pre-symptomatic D5 stage. Shaded areas indicate the group distribution. **(C)** Unsupervised hierarchical clustering heatmap of normalized protein-abundance profiles from three CTRL and three ERO1L-OE biological replicates. Rows represent quantified proteins and columns represent individual head samples. Color indicates relative protein abundance expressed as row-scaled Z-scores, from low (blue) to high (red). The annotation bar identifies CTRL and ERO1L-OE samples.

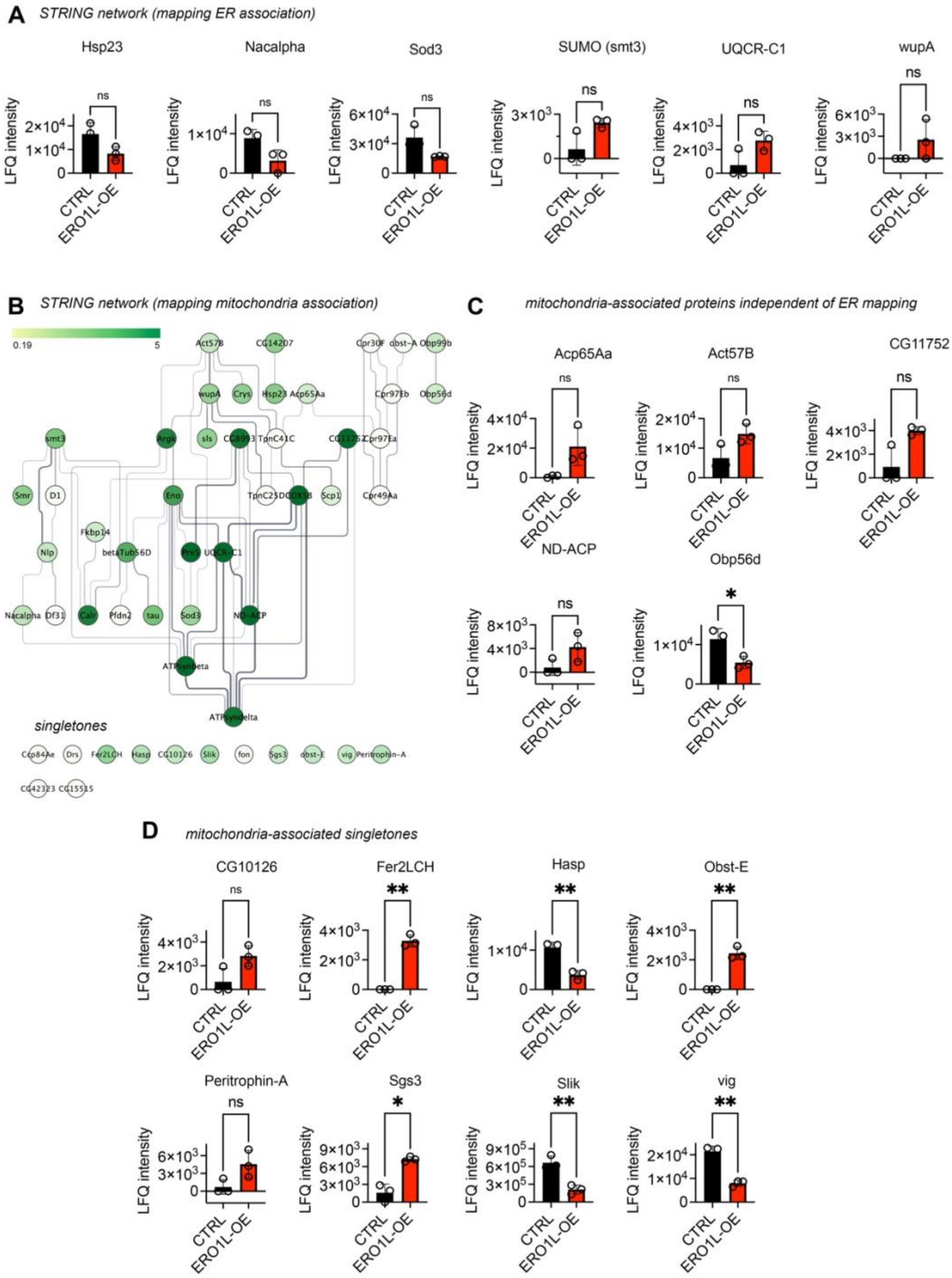


**Supplementary Figure 2 Selective remodeling of ER- and mitochondria-associated protein networks in presymptomatic neuronal *ERO1L* expressing heads.**

**(A)** LFQ intensity profiles of proteins mapped to the ER-associated STRING network that were not significantly altered in 5-day-old *nSyb>ERO1L* (ERO1L-OE) heads relative to controls: heat-shock protein 23 (Hsp23), N-α-acetyltransferase alpha subunit (Ncalpha), extracellular superoxide dismutase 3 (Sod3), SUMO (smt3), ubiquinol–cytochrome *c* reductase core protein 1 (UQCR-C1), and Troponin I/wing up

A (wupA). **(B)** STRING functional-association network of mitochondria-associated proteins identified in the proteomic dataset. The network was visualized in Cytoscape using the yFiles Orthogonal Layout. Node color represents the mitochondria-association mapping score, with lighter to darker green indicating lower to higher scores, respectively. Edges represent STRING functional associations. Proteins not connected to the main network are shown as singletons. **(C)** LFQ intensity profiles of mitochondria-associated proteins identified independently of ER mapping: adult cuticle protein 65Aa (Acp65Aa), actin 57B (Act57B), CG11752, NADH dehydrogenase subunit ACP (ND-ACP), and odorant-binding protein 56d (Obp56d). Of these proteins, only Obp56d was significantly reduced in *nSyb>ERO1L* heads. **(D)** LFQ intensity profiles of mitochondria-associated singleton proteins: CG10126, ferritin 2 light-chain homolog (Fer2LCH), Hasp, Obstructor-E (Obst-E), Peritrophin-A, salivary gland secretion 3 (Sgs3), Slingshot (Slik), and Vasa intronic gene (vig). Fer2LCH, Obst-E, and Sgs3 were increased, whereas Hasp, Slik, and vig were decreased in *nSyb>ERO1L* heads. CG10126 and Peritrophin-A were not significantly altered. Circles represent individual biological replicates. Statistical comparisons of LFQ intensities were performed using unpaired two-tailed *t*-tests with Welch's correction. ns, not significant; **P* < 0.05; ***P* < 0.01. Together, these data indicate that neuronal *ERO1L* overexpression at the presymptomatic stage selectively remodels subsets of ER- and mitochondria-associated proteins.

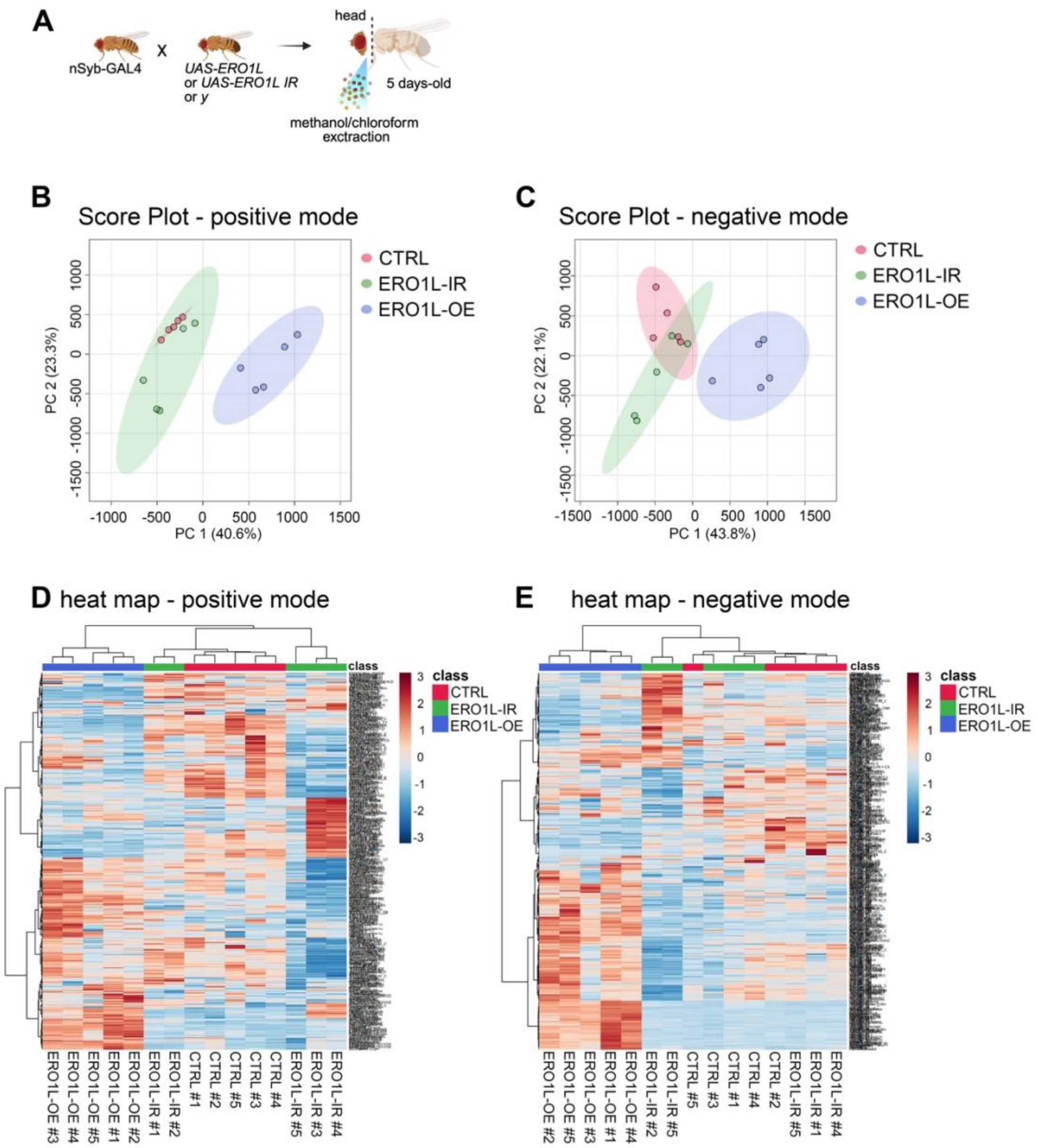


**Supplementary Figure 3 Global metabolomic profiles of control, neuronal *ERO1L*-inhibited, and neuronal *ERO1L*-overexpressing fly heads at the pre-symptomatic stage.**

**(A)** Experimental workflow for metabolomic profiling. nSyb-GAL4 flies were crossed to *UAS-ERO1L* to generate neuronal ERO1L-overexpressing flies (ERO1L-OE), to the *ERO1L* RNAi line e (ERO1L-IR), or to the y control line (CTRL). Heads were collected from 5-day-old adults and metabolites were extracted using methanol/chloroform. (**B–C)** Principal-component analysis score plots of metabolomic profiles acquired in positive-ion mode **(B)** and negative-ion mode **(C)**. Each point represents one biological replicate. CTRL, ERO1L-IR, and ERO1L-OE samples show distinct group distributions in both acquisition modes. In positive-ion mode, principal component 1 (PC1) and principal component 2 (PC2) explain 40.6% and 23.3% of the variance, respectively.

In negative-ion mode, PC1 and PC2 explain 43.8% and 22.1% of the variance, respectively. Shaded ellipses indicate group distributions. **(D–E)** Unsupervised hierarchical clustering heatmaps of metabolite-abundance profiles acquired in positive-ion mode **(D)** and negative-ion mode **(E)**. Columns represent individual biological replicates from CTRL, ERO1L-IR, and ERO1L-OE groups; rows represent detected metabolite features. Color indicates relative abundance expressed as row-scaled Z-scores, from low (blue) to high (red). The annotation bar identifies the experimental group for each sample.

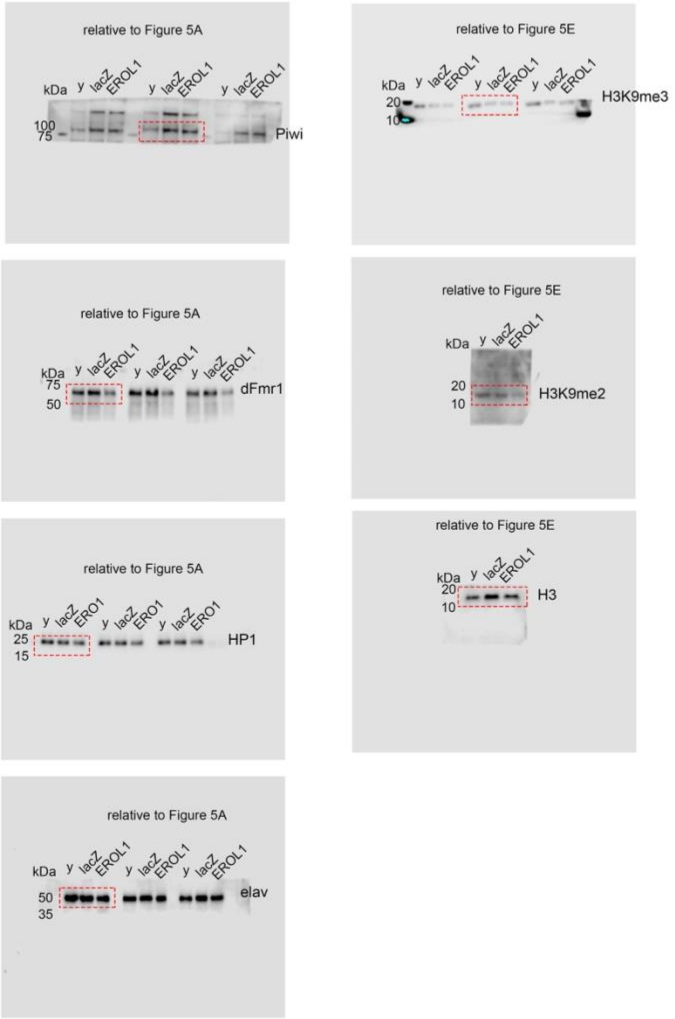


**Supplementary Figure 4** Uncropped western blot images supporting quantification shown in Figure 5. Red boxes indicate the cropped areas used for densitometric quantification

**Suppplementary Table 1 List of primers used in this study**

| Target | Forward primer (5′–3′) | Reverse primer (5′–3′) |
|---|---|---|
| *297* | GGTGATCCAGAAACCCTTCA | CTTTCGATGGCTCCCAGTAG |
| *412* | CACCGGTTTGGTCGAAAG | GGACATGCCTGGTATTTTGG |
| *Act5C* | CAACTGGGACGATATGGAGAAG | GTCTCGAACATGATCTGGGTC |
| *Blood* | TATCGCATGGCAGATAGCCAAA | CGTGGAATTCGGAAGTGGTTTC |
| *Copia* | GCATGAGAGGTTTGGCCATATAAGC | GGCCCACAGACATCTGAGTGTACTACA |
| *ERO1L* | GTGCAGAGAAATCTGTGGGC | TTTTTGCTAGTCTCCGTCTCATC |
| *GAL4* | TTGAAATCGCGTCGAAGGA | GGCTCCAATGGCTAATATGCA |
| *Gypsy12* | TGACTCGGCTGATGTTTCTC | GAAACACAGGTGGAATCGTG |
| *RpL11* | CCATCGGTATCTATGGTCTGGA | CATCGTATTTCTGCTGGAACCA |
| *RpL32* | CATCCGCCCAGCATACAG | CCATTTGTGCGACAGCTTAG |
| *RP49* | TCCTACCAGCTTCAAGATGACC | CACGTTGTGCACCAGGAACT |
| *Roo* | CGTCTGCAATGTACTGGCTCT | CGGCACTCCACTAACTTCTCC |
| *Tabor* | GAGCAAGAATTATGCTCGAAGAA | AATTTATGTCCGGTTTCGTTTTT |

**Supplementary Table 2 Dyes, probes, and antibodies used in this study**

| Reagent / antibody | Target / description | Host / type | Supplier | Catalogue number / clone |
|---|---|---|---|---|
| DAPI | Nuclear DNA counterstain; 4′,6-diamidino-2-phenylindole dihydrochloride | Fluorescent DNA stain | Merck | 10236276001 |
| DHE | Cell-permeable redox-sensitive fluorescent probe used to assess superoxide-associated ROS fluorescence | Fluorescent chemical probe. | Merck | 309800 |
| Alexa Fluor 488 Phalloidin | F-actin | Fluorescent probe | Thermo Fisher Scientific | A12379 |
| Anti-HP1 | Heterochromatin Protein 1 (HP1) | Mouse monoclonal | DSHB | C1A9 |
| Anti-H3K9me2 | Histone H3 lysine 9 dimethylation (H3K9me2) | Rabbit polyclonal | Invitrogen | PA5-16195 |
| Anti-H3K9me3 | Histone H3 lysine 9 trimethylation (H3K9me3) | Rabbit polyclonal | Invitrogen | 600-401-I71 |
| Anti-Histone H3 | Histone H3 | Rabbit monoclonal | Cell Signaling Technology | 9717 |
| Anti-dFmr1 | Drosophila fragile X mental retardation protein 1 (dFmr1) | Mouse monoclonal | DSHB | 5A11 |
| Anti-Piwi | Piwi | Rabbit polyclonal | Abcam | ab5207 |
| Anti-Elav | Embryonic lethal abnormal vision (Elav) | Mouse monoclonal | DSHB | 9F8A9 |
| Goat anti-mouse IgG-HRP | Mouse primary antibodies | HRP-conjugated secondary antibody | Invitrogen | 31430 |
| Goat anti-rabbit IgG-HRP | Rabbit primary antibodies | HRP-conjugated secondary antibody | Invitrogen | 31460 |
| Goat anti-mouse IgG, Alexa Fluor 594 | Mouse primary antibodies | Alexa Fluor 594-conjugated secondary antibody | Invitrogen | A-11032 |

**Supplementary Table 3 Differentially abundant proteins identified in ERO1L-OE versus CTRL *Drosophila melanogaster* heads**

| Protein | $\log_2$ fold change | Raw P value |
|---|---|---|
| ATPsyndelta | 4.4496 | 0.012189 |
| Crys | 4.3722 | 0.010865 |
| Acp65Aa | 4.2644 | 0.053178 |
| CG8369 | 4.1059 | 0.010757 |
| sls | 4.0866 | 0.076704 |
| wupA | 4.0718 | 0.003275 |
| ATPsynbeta | 3.8458 | 0.011175 |
| TpnC25D | 3.4995 | 0.001477 |
| betaTub56D | 3.4179 | 0.002778 |
| Calr | 3.4026 | 0.004846 |
| Fer2LCH | 3.3403 | 0.000153 |
| Argk1 | 3.3167 | 0.002004 |
| CG12990 | 3.2079 | 0.001216 |
| obst-E | 2.9151 | 0.000572 |
| COX5B | 2.8683 | 0.000483 |
| CG16713 | 2.5411 | 0.008910 |
| RpLP2 | 2.4719 | 0.063498 |
| TotA | 2.4664 | 0.042980 |
| DptA | 2.3522 | 0.075498 |
| Ref1 | 2.3067 | 0.050106 |
| Peritrophin-A | 2.2977 | 0.064824 |
| CG31705 | 2.2612 | 0.002616 |
| Sgs3 | 2.0919 | 0.002923 |
| ND-ACP | 2.0893 | 0.098149 |
| CG8993 | 2.0720 | 0.021472 |
| Drs | 2.0662 | 0.000595 |
| Nlp | 2.0647 | 0.037085 |
| Eno | 2.0279 | 0.007705 |
| Cpr97Eb | 1.8725 | 0.024125 |
| Scp1 | 1.8318 | 0.000692 |
| CG11752 | 1.7954 | 0.032689 |
| CG7781 | 1.7332 | 0.001055 |
| CG10126 | 1.7080 | 0.056537 |
| CG15515 | 1.6891 | 0.004499 |
| UQCR-C1 | 1.6023 | 0.068280 |
| TpnC41C | 1.6003 | 0.013040 |
| Cpr49Aa | 1.5649 | 0.008727 |
| CG42367 | 1.5372 | 0.000508 |
| Sumo | 1.5205 | 0.051532 |
| Pfdn2 | 1.4892 | 0.083221 |
| CG42337 | 1.4620 | 0.051593 |
| Cbp53E | 1.3659 | 0.008521 |
| Prx5 | 1.3525 | 0.010074 |

| Protein | $\log_2$ fold change | Raw P value |
|---|---|---|
| Cpr30F | 1.3055 | 0.045871 |
| CG14207 | 1.1980 | 0.009507 |
| Act57B | 1.1617 | 0.057942 |
| Glos | 1.1398 | 0.038170 |
| LKRSDH | 1.0272 | 0.016114 |
| Hsp23 | −1.0033 | 0.062664 |
| Obp99b | −1.0189 | 0.033773 |
| Obp56d | −1.0637 | 0.028806 |
| Acbp2 | −1.0863 | 0.015300 |
| Sod3 | −1.1001 | 0.043407 |
| Fkbp14 | −1.1053 | 0.001926 |
| CG7203 | −1.1513 | 0.059304 |
| Smr | −1.1993 | 0.014947 |
| Cpr97Ea | −1.2067 | 0.004582 |
| Ccp84Ae | −1.2091 | 0.096160 |
| CG44242 | −1.2296 | 0.005293 |
| Akap200 | −1.2545 | 0.005713 |
| vig | −1.4279 | 0.000780 |
| Nacalpha | −1.4376 | 0.047981 |
| D1 | −1.4729 | 0.008510 |
| tau | −1.4921 | 0.002165 |
| fon | −1.5076 | 0.005812 |
| CG42323 | −1.5195 | 0.081653 |
| obst-A | −1.5234 | 0.019191 |
| Hasp | −1.5321 | 0.002316 |
| CG7214 | −1.5422 | 0.026554 |
| Slik | −1.6605 | 0.004355 |
| l(2)03659 | −1.7366 | 0.006938 |
| Df31 | −1.7575 | 0.043685 |
| CG14764 | −1.8971 | 0.020961 |
| ND-AGGG | −2.1378 | 0.031633 |
| trp | −2.1441 | 0.008142 |
| Sem1 | −2.1992 | 0.028708 |
| RpS30 | −4.0228 | 0.015909 |

**Supplementary Table 4 Variable importance in projection (VIP) scores for metabolites discriminating ERO1L-OE and CTRL *Drosophila melanogast*er heads in positive ionization**

| Metabolite | VIP score (Component 1) |
|---|---|
| C15H21N5O8; PlaSMA ID-968.2 | 6.9384 |
| C15H21N5O8; PlaSMA ID-968.1 | 5.2576 |
| 1-isothiocyanato-8-methanesulfinyloctane; PlaSMA ID-931 | 4.4171 |
| Phosphocholine.1 | 4.2664 |
| Pesticide5_Carboxin_C12H13NO2S_Vitavax | 3.6760 |
| haplamine | 3.3793 |
| Hypoxanthine.4 | 3.3044 |
| C15H21N5O8; PlaSMA ID-968 | 3.2679 |
| Hypoxanthine.2 | 3.2559 |
| O-Phosphocolamine | 3.2090 |
| Deoxyadenosine; PlaSMA ID-1068 | 2.9390 |
| Phosphocholine | 2.5557 |
| Hypoxanthine | 2.5459 |
| Hypoxanthine.3 | 2.4618 |
| 1-isothiocyanato-7-methanesulfinylheptane; PlaSMA ID-828 | 2.3763 |
| NCGC00380822-01!1-[(Z)-but-2-en-2-yl]-8-chloro-3,9-dihydroxy-4,7-dimethylbenzo[b][1,4]benzodioxepin-6-one | 2.3614 |
| INDOLEACETALDEHYDE | 2.0871 |
| INDOLEACETALDEHYDE.1 | 2.0751 |
| 1-(pyrimidin-2-yl)pyrrolidine-2-carboxylic acid | 2.0495 |
| narcotine | 2.0435 |
| 2-Thiocytidine.1 | 2.0364 |
| Zafirlukast | 1.9887 |
| Shikimic Acid | 1.9137 |
| Euphorbiasteroid | 1.8278 |
| Pantothenic acid | 1.8025 |
| (R)-2-amino-3-(1H-indol-3-yl)-N-methylpropanamide oxalate | 1.7903 |
| Hypoxanthine.1 | 1.6882 |
| (Z)-2-(4-(methoxycarbonyl)benzylidene)-7-(morpholino-4-iummethyl)-3-oxo-2,3-dihydrobenzofuran-6-olate | 1.6795 |
| (2S,3S)-3-hydroxy-2-((S)-2-((S)-2-(2-hydroxy-4-oxoquinazolin-3(4H)-yl)-3-(1H-indol-3-yl)propanamido)propanamido)butanoic acid | 1.6474 |
| 2-Thiocytidine | 1.5830 |
| Unknown (carbon number 11); PlaSMA ID-1285 | 1.5702 |
| N-2-Fluorenylacetamide | 1.5372 |
| Acetaminophen glucuronide | 1.5230 |
| Unknown (carbon number 6); PlaSMA ID-176 | 1.5008 |
| Ononin; PlaSMA ID-2341 | 1.4908 |
| Remerine; PlaSMA ID-1342 | 1.4839 |
| NCGC00384999-01_C24H24O10_2-({6-O-[(2E)-3-(4-Hydroxyphenyl)-2-propenoyl]-beta-D-glucopyranosyl}oxy)-3-phenylacrylic acid | 1.4694 |
| Unknown (carbon number 11); PlaSMA ID-1180 | 1.4302 |

| Metabolite | VIP score (Component 1) |
|---|---|
| salviaflaside | 1.4241 |
| Curcumin | 1.4223 |
| Ganoderic Acid F.1 | 1.4059 |
| Tolfenamic acid | 1.3430 |
| NCGC00384929-01!(2S,3S)-3,5,7-trihydroxy-2-[4-hydroxy-3-[(2S,3R,4S,5S,6R)-3,4,5-trihydroxy-6-(hydroxymethyl)oxan-2-yl]oxyphenyl]-2,3-dihydrochromen-4-one | 1.3124 |
| Sulfamethoxazole | 1.3098 |
| NCGC00384999-01_C24H24O10_2-({6-O-[(2E)-3-(4-Hydroxyphenyl)-2-propenoyl]-beta-D-glucopyranosyl}oxy)-3-phenylacrylic acid.1 | 1.3028 |
| Acacetin-7-O-rutinoside; PlaSMA ID-2971 | 1.2769 |
| 1-isothiocyanato-7-methanesulfinylheptane; PlaSMA ID-828.1 | 1.2016 |
| NCGC00169840-02![2-(4-hydroxy-3-methoxyphenyl)-4-[(4-hydroxy-3-methoxyphenyl)methyl]oxolan-3-yl]methyl (E)-3-(4-hydroxyphenyl)prop-2-enoate | 1.2014 |
| 1-Isothiocyanato-9-(methylsulfinyl)-nonane; LC-ESI-QTOF; MS2; CE | 1.2007 |
| Biuret | 1.1617 |
| Cyanidin | 1.1461 |
| Ganoderic Acid F | 1.1417 |
| Unknown (carbon number 7); PlaSMA ID-718 | 1.1251 |
| N-[1-(4-methoxy-6-oxopyran-2-yl)-2-methylbutyl]acetamide.3 | 1.1121 |
| N-Acetylarginine; CE0; SNEIUMQYRCDYCH-LURJTMIESA-N | 1.0676 |
| inosine.3 | 1.0632 |
| 3-[(E)-2-(3-hydroxyphenyl)ethenyl]-5-methoxyphenol.1 | 1.0567 |
| Di-dihydrocaffeoyl spermidine; PlaSMA ID-2573 | 1.0567 |
| Unknown (carbon number 16); PlaSMA ID-1196.1 | 1.0506 |
| 4-hydroxy-3-[(4-hydroxy-6-methyl-2-oxopyran-3-yl)methyl]-6-methylpyran-2-one | 1.0473 |
| N-Fructosyl isoleucine; PlaSMA ID-1287.3 | 1.0377 |
| N-Fructosyl isoleucine; PlaSMA ID-1287 | 1.0344 |
| NCGC00385338-01!4-[3,4,5-trihydroxy-6-(hydroxymethyl)oxan-2-yl]oxy-9-(3,4,5-trimethoxyphenyl)-5a,6,8a,9-tetrahydro-5H-[2]benzofuro[6,5-f][1,3]benzodioxol-8-one | 1.0182 |
| Unknown (carbon number 11); PlaSMA ID-199 | 1.0146 |
| N-Fructosyl isoleucine; PlaSMA ID-1287.1 | 1.0139 |
| C15H19NO6; PlaSMA ID-1635 | 1.0084 |
| INDOLEACETALDEHYDE.3 | 1.0075 |
| Gluconolactone | 1.0045 |

**Supplementary Table 5 Variable importance in projection (VIP) scores for metabolites discriminating ERO1L-OE and CTRL *Drosophila melanogast*er heads in negative ionization**

| Metabolite name | VIP score (Component 1) |
|---|---|
| Gluconic acid | 5.7379 |
| Unknown (carbon number 19); PlaSMA ID-751 | 5.2035 |
| Unknown.20 | 4.3501 |
| Unknown.17 | 4.0777 |
| HippuricAcid | 3.3403 |
| HippuricAcid.1 | 3.3403 |
| (1R,2E,8S,10R,11S)-10,11-dihydroxy-6-(methoxymethyl)-1,10-dimethyl-5-oxo-4,14-dioxatricyclo[9.2.1.0?,?]tetradeca-2,6-dien-8-yl acetate | 2.8961 |
| ALPHA-KETOGLUTARIC ACID | 2.8146 |
| Ascorbic acid | 2.8129 |
| Unknown.13 | 2.6673 |
| LPE 18:1; PlaSMA ID-1334 | 2.6482 |
| Unknown.79 | 2.6357 |
| Glutamic acid | 2.5907 |
| NCGC00380982-01_C22H25NO8_8-Benzoyl-2-[(3E)-1,2-dihydroxy-3-hexen-1-yl]-9-hydroxy-8-methoxy-3-methyl-1-oxa-7-azaspiro[4.4]non-2-ene-4,6-dione | 2.5688 |
| Unknown.206 | 2.4532 |
| Unknown.208 | 2.4490 |
| N-Acetylmannosamine | 2.4242 |
| Unknown.19 | 2.4003 |
| NCGC00386071-01_[3-methyl-1-[3-methyl-1-oxo-1-(2,3,4,5,6-pentahydroxyhexoxy)pentan-2-yl]oxy-1-oxopentan-2-yl] 2-hydroxy-3-methylpentanoate | 2.3960 |
| 5-Methoxypsoralen | 2.3616 |
| Unknown.166 | 2.3389 |
| Unknown.90 | 2.2981 |
| LPE 18:2; PlaSMA ID-1316 | 2.2046 |
| Maltotriose | 2.2046 |
| Unknown.21 | 2.1954 |
| Unknown.190 | 2.1574 |
| Fenoldopam.1 | 2.1098 |
| O-Phosphocolamine | 2.1014 |
| NCGC00347571-02_C27H40O4_(3beta,5alpha,8xi,14xi,25S)-3-Hydroxyspirost-9(11)-en-12-one | 2.0363 |
| 7-Methylsulfinylheptyl isothiocyanate; PlaSMA ID-311 | 2.0183 |
| Unknown.24 | 2.0046 |
| Flavone base + 4O, 1Prenyl; PlaSMA ID-835 | 2.0043 |
| 2-Oxobutyric acid | 1.9883 |
| Unknown.98 | 1.9682 |
| FT-thioether; C10H18F3NO4S2 | 1.9502 |
| INOSINE.2 | 1.9043 |

| Metabolite name | VIP score (Component 1) |
| --- | --- |
| Unknown.23 | 1.8919 |
| Unknown (carbon number 5); PlaSMA ID-88.1 | 1.8501 |
| Unknown.18 | 1.8113 |
| Fenoldopam.3 | 1.7823 |
| Tenofovir | 1.7653 |
| NCGC00180624-03_C25H30O13_2-Propenoic acid, 3-(3-hydroxy-4-methoxyphenyl)-, (1aS,1bS,2S,5aR,6S,6aS)-2-(beta-D-glucopyranosyloxy)-1a,1b,2,5a,6,6a-hexahydro-1a-(hydroxymethyl)oxireno[4,5]cyclopenta[1,2-c]pyran-6-yl ester, (2E)- | 1.7600 |
| Pantothenic acid (Not validated); PlaSMA ID-312 | 1.7479 |
| Unknown.51 | 1.7468 |
| Unknown.182 | 1.7160 |
| NCGC00384589-01_[(2R,3R,4S,5R,6R)-6-[2-(3,4-dihydroxyphenyl)ethoxy]-3,5-dihydroxy-4-[(2S,3R,4R,5R,6S)-3,4,5-trihydroxy-6-methyloxan-2-yl]oxyoxan-2-yl]methyl (E)-3-(4-hydroxy-3-methoxyphenyl)prop-2-enoate | 1.7012 |
| Succinic acid | 1.6898 |
| Unknown.7 | 1.6771 |
| Fenoldopam | 1.6688 |
| (2R)-5-methoxy-2-methyl-2,3,8,9-tetrahydrofuro[2,3-h]chromen-4-one | 1.6155 |
| Unknown.115 | 1.5829 |
| 2,8-Quinolinediol | 1.5795 |
| Unknown.6 | 1.5784 |
| NCGC00380467-01_C26H30O13_2H-Pyran-4-acetic acid, 5-carboxy-2-(beta-D-glucopyranosyloxy)-3,4-dihydro-3-[2-[[(2Z)-1-oxo-3-phenyl-2-propen-1-yl]oxy]ethylidene]-, alpha-methyl ester, (2S,3Z,4S)- | 1.5784 |
| XANTHURENIC ACID | 1.5708 |
| INOSINE | 1.5622 |
| Unknown.202 | 1.5606 |
| Fenoldopam.4 | 1.5186 |
| C13H17NO2S; PlaSMA ID-394 | 1.5182 |
| Dihydrostilbene base + 3O, 1carboxy, O-Hex; PlaSMA ID-1149 | 1.5006 |
| INOSINE.1 | 1.4807 |
| Fenoldopam.2 | 1.4766 |
| Hypoxanthine | 1.4704 |
| Unknown.180 | 1.4402 |
| Unknown.76 | 1.4287 |
| Maltotriose; LC-ESI-QTOF; MS2; CE | 1.4072 |
| Unknown.138 | 1.3818 |
| MBOA; PlaSMA ID-134 | 1.3575 |
| 2-Chloro-N6-cyclopentyladenosine.1 | 1.3554 |
| Unknown.86 | 1.3469 |
| NCGC00380230-01_C26H40O9_2,4a,6-Trihydroxy-8-isopropyl-1,1-dimethyl-2,3,4,4a,5,10,11,11a-octahydro-1H-dibenzo[a,d][7]annulen-7-yl beta-D-glucopyranoside | 1.3417 |
| Unknown.167 | 1.3413 |
| FT-OH; C3H5F3O | 1.3174 |

| Metabolite name | VIP score (Component 1) |
| --- | --- |
| Unknown.136 | 1.2743 |
| Atropine | 1.2616 |
| 2-Chloro-N6-cyclopentyladenosine | 1.2563 |
| Unknown.89 | 1.2418 |
| Grossamide or its isomer (Not validated); PlaSMA ID-1722 | 1.2359 |
| Unknown.39 | 1.2315 |
| Unknown.75 | 1.2282 |
| XANTHINE | 1.2213 |
| Unknown.33 | 1.2192 |
| Chloridazon | 1.2181 |
| C23H42NO7P; PlaSMA ID-1303 | 1.2165 |
| Unknown.66 | 1.2052 |
| 2-Ketogluconic acid.1 | 1.2006 |
| Unknown.63 | 1.1961 |
| Formononetin-7-O-glucoside; PlaSMA ID-1106 | 1.1922 |
| Unknown.2 | 1.1849 |
| Unknown.56 | 1.1799 |
| N-Fructosyl pyroglutamate; PlaSMA ID-534 | 1.1792 |
| Xanthosine (Not validated); PlaSMA ID-511 | 1.1788 |
| Unknown.142 | 1.1603 |
| Raltitrexed | 1.1588 |
| Unknown.150 | 1.1511 |
| Inosine-5-monophosphate | 1.1476 |
| N-2-Fluorenylacetamide | 1.1388 |
| Unknown.193 | 1.1270 |
| 3-Nitrotyrosine | 1.1263 |
| NCGC00169614-02_C24H21N5O4_(1S,3S,12S,14R,27S)-12-Hydroxy-1,3-dimethyl-2,5,15,23,25-pentaazaheptacyclo[12.10.2.1~2,5~.0~6,11~.0~12,27~.0~15,24~.0~17,22~]heptacosa-6,8,10,17,19,21,23-heptaene-4,16,26-trione | 1.1176 |
| Unknown.157 | 1.1084 |
| NCGC00385087-01_3-[5,7-dihydroxy-2-(4-methoxyphenyl)-4-oxo-2,3-dihydrochromen-3-yl]-5,7-dihydroxy-2-(4-hydroxyphenyl)-2,3-dihydrochromen-4-one | 1.0843 |
| Unknown.71 | 1.0842 |
| Unknown.9 | 1.0690 |
| 2-Ketogluconic acid | 1.0664 |
| (3aR,4R,6E,10Z,11aR)-4-hydroxy-10-(hydroxymethyl)-3-methylidene-2-oxo-3a,4,5,8,9,11a-hexahydrocyclodeca[b]furan-6-carbaldehyde | 1.0643 |
| Unknown (carbon number 19); PlaSMA ID-751.3 | 1.0619 |
| Omeprazole | 1.0535 |
| NCGC00381006-01_C20H26O10_2-(beta-D-Glucopyranosyloxy)benzyl 1,6-dihydroxy-2-cyclohexene-1-carboxylate | 1.0509 |
| Unknown.35 | 1.0294 |
| Unknown (carbon number 19); PlaSMA ID-751.2 | 1.0258 |

### Supplementary Table 6 Abundance of KEGG-mapped metabolites in controls and ERO1L-OE *Drosophila melanogaster* samples

| Metabolite name | CTRL1 | CTRL2 | CTRL3 | CTRL4 | CTRL5 | ERO_IR1 | ERO_IR2 | ERO_IR3 | ERO_IR4 | ERO_IR5 | ERO_OE1 | ERO_OE2 | ERO_OE3 | ERO_OE4 | ERO_OE5 |
|---|---|---|---|---|---|---|---|---|---|---|---|---|---|---|---|
| Phosphorylcholine | 180681.7 | 167306.5 | 220103.3 | 187476.1 | 169951.1 | 184418.4 | 192312.3 | 222028.1 | 212839.3 | 151236.3 | 227191.7 | 200689 | 232999.2 | 234896 | 227460.6 |
| Hypoxanthine | 41808.13 | 48370.83 | 36660.61 | 35392.04 | 19964.12 | 43519.17 | 40151.91 | 49764.61 | 40419.85 | 22421.97 | 47284.95 | 44810.72 | 75418.8 | 81848.33 | 60990.5 |
| O-Phosphoethanolamine | 12469.97 | 15618.61 | 13467.25 | 13188.5 | 16183.62 | 15276.31 | 25102.9 | 15776.1 | 16189 | 24349.57 | 25087.41 | 17662.93 | 25831.18 | 25601.8 | 17853.8 |
| Indoleacetaldehyde | 4239.385 | 1063.198 | 1837.249 | 2789.082 | 5271.098 | 5205.772 | 4730.862 | 1591.802 | 2615.459 | 3241.171 | 5735.228 | 8352.171 | 3718.354 | 2677.056 | 1511.354 |
| (-)-alpha-Narcotine | 16303.91 | 15806.05 | 15484.73 | 15635.54 | 18617.43 | 18231.69 | 19057.89 | 8443.026 | 8070.106 | 9907.124 | 9160.376 | 7946.852 | 10333.8 | 9268.777 | 8602.083 |
| Zafirlukast | 16976.37 | 15749.98 | 13866.49 | 14154.25 | 16320.56 | 13445.82 | 14191.7 | 6815.921 | 6840.257 | 9431.424 | 8550.432 | 8440.922 | 8726.583 | 7993.525 | 8880.245 |
| Shikimic acid | 4776.928 | 5439.873 | 4181.84 | 4280.757 | 5345.387 | 6074.136 | 5780.249 | 5801.623 | 5917.363 | 4353.75 | 12466.42 | 12227.73 | 10847.83 | 10961.57 | 9500.274 |
| Pantothenic acid | 8290.584 | 7891.07 | 9122.727 | 8923.058 | 9918.239 | 8904.523 | 8600.088 | 13827.04 | 13503.31 | 7565.537 | 14808.81 | 14828.61 | 14034.38 | 13189.34 | 15737.59 |
| 2-Acetylaminofluorene | 8647.025 | 8236.22 | 8339.773 | 9508.4 | 9283.392 | 6207.424 | 6639.634 | 8785.259 | 9388.608 | 6250.651 | 4983.369 | 4814.385 | 4530.997 | 4532.702 | 4850.808 |
| Acetaminophen glucuronide | 15981.54 | 14793.17 | 17907.9 | 17333.35 | 19043.54 | 16972.82 | 17330.21 | 30310.86 | 30735.98 | 19062.63 | 26285.9 | 26467.92 | 19231.34 | 17334.41 | 23615.03 |
| Salviaflaside | 6932.109 | 6488.341 | 5606.235 | 5200.663 | 4106.838 | 4690.61 | 5282.34 | 5134.041 | 4969.078 | 2861.477 | 2068.106 | 1838.978 | 2131.361 | 1917.852 | 2028.379 |
| Curcumin | 5867.417 | 5507.986 | 6036.877 | 6447.595 | 7134.852 | 4467.7 | 5014.329 | 4686.566 | 4572.075 | 3415.282 | 2849.059 | 2559.75 | 2753.496 | 2800.699 | 2497.206 |
| Tolfenamic acid | 5035.003 | 5598.086 | 6294.537 | 6728.841 | 7073.45 | 4336.846 | 4301.793 | 6809.013 | 6966.862 | 4298.908 | 2993.594 | 2874.738 | 2658.588 | 2648.376 | 3425.77 |
| Sulfamethoxazole | 16074.34 | 11308.74 | 25164.21 | 24358.24 | 20190.44 | 22967.49 | 22418.45 | 16894.82 | 17138.54 | 15858.06 | 11335.11 | 8354.857 | 11560.48 | 18361.87 | 19185.67 |
| Biuret | 4331.993 | 3562.893 | 4304.889 | 5580.367 | 4906.535 | 5313.79 | 5927.016 | 4823.09 | 5595.545 | 3617.809 | 1660.296 | 1830.345 | 2004.022 | 2911.151 | 1744.968 |
| Cyanidin | 20567.58 | 20381.62 | 18492.82 | 18143.48 | 17109.62 | 15428.26 | 16919.2 | 12836.66 | 12882.68 | 12810.07 | 16110.98 | 14773.97 | 16654.52 | 16166.57 | 16971.08 |
| Ganoderic acid F | 1096.591 | 1093.349 | 1435.147 | 968.3135 | 807.8964 | 857.1831 | 655.6591 | 230.7501 | 352.7167 | 305.9726 | 710.7915 | 770.3306 | 1460.338 | 1596.697 | 1157.507 |
| Inosine | 10605.36 | 11427.21 | 7355.774 | 10487.66 | 11147.81 | 13722.93 | 11975.7 | 9592.62 | 13586 | 9650.461 | 17209.43 | 15244.14 | 12608.58 | 17795.8 | 22547.5 |
| Gluconolactone | 2025.221 | 1871.876 | 1547.356 | 1718.416 | 1134.998 | 1068.681 | 1110.791 | 915.2617 | 862.483 | 1042.044 | 3663.65 | 3195.16 | 3478.748 | 3464.789 | 3332.561 |
| Gluconic acid | 43958.62 | 40766.56 | 26205.41 | 53296.11 | 44032.46 | 22514.1 | 23397.58 | 25427.04 | 26886.83 | 24393.65 | 108927.2 | 103036.8 | 72842.39 | 99260.32 | 94344.63 |
| Hippuric acid | 113112.2 | 54355.06 | 46394.07 | 11764.8 | 13931.4 | 31777.48 | 13091.56 | 21440.79 | 13451.86 | 5966.249 | 18792.48 | 19399.05 | 12153.08 | 13109.37 | 14547.83 |

| | | | | | | | | | | | | | | | |
|---|---|---|---|---|---|---|---|---|---|---|---|---|---|---|---|
| Oxoglutaric acid (a-KG) | 10306.64 | 14316.9 | 9378.519 | 10646.31 | 14839.96 | 16948.84 | 4886.46 | 12849.8 | 18495.4 | 4334.367 | 21062.97 | 26946.68 | 26821.47 | 20556.55 | 31200.8 |
| Ascorbic acid | 11855.05 | 15698.35 | 12598.32 | 12581.5 | 14761.82 | 17602.2 | 2526.091 | 11908.71 | 18549.45 | 2139.974 | 21588.82 | 34330.77 | 26209.85 | 20686.67 | 37149.03 |
| Glutamic acid | 22834.27 | 30559.41 | 21234.23 | 26325.35 | 30912.03 | 30795.9 | 46497 | 26122.1 | 35607.5 | 42836.5 | 41712.42 | 36438.36 | 37385.55 | 40912.29 | 34430.5 |
| N-Acetylmannosamine | 7019.021 | 5664.567 | 4610.796 | 6030.68 | 5070.34 | 6184.626 | 3387.652 | 4037.443 | 5683.915 | 2815.375 | 16486.09 | 15438.01 | 12168.49 | 17216.55 | 13897.66 |
| Bergapten | 63218.89 | 72844.22 | 65795.69 | 88446.08 | 96322.5 | 59796.4 | 81988.65 | 78543.65 | 84545.31 | 83359.14 | 103744.6 | 108614.9 | 72443.5 | 92990.33 | 88935.92 |
| Maltotriose | 59217.35 | 62465.49 | 57258.22 | 63935.44 | 66382.14 | 55079.98 | 28910.3 | 45456.68 | 61321.32 | 28249.98 | 79303.21 | 85290.09 | 55135.63 | 69237.35 | 81411.67 |
| Fenoldopam | 4733.95 | 4717.467 | 3769.326 | 4370.55 | 3554.987 | 4014.154 | 3961.429 | 3146.074 | 3500.039 | 3588.349 | 12726.78 | 9411.761 | 7290.506 | 9758.676 | 6849.529 |
| 2-Ketobutyric acid (2-KB) | 6249.15 | 9010.189 | 5236.365 | 6815.337 | 8653.859 | 9602.548 | 2692.69 | 7794.01 | 10795.3 | 2416.171 | 11736.57 | 15486.42 | 15307.09 | 10544.59 | 18711.7 |
| Tenofovir | 26631.07 | 25382.59 | 23012.07 | 30062 | 26120.43 | 20304.81 | 22543.64 | 20190.57 | 22775.96 | 21222.58 | 21172.88 | 19574.04 | 22686.16 | 20607.07 | 18700.85 |
| Succinic acid | 20257.47 | 20321.71 | 9749.696 | 27162.29 | 21674.18 | 20729.35 | 20052.6 | 7745.03 | 23565.4 | 22887.09 | 31644.23 | 32295.99 | 16054.87 | 28313.35 | 29875.7 |
| Xanthurenic acid | 2893.285 | 3370.77 | 2773.451 | 2951.332 | 3039.586 | 3208.763 | 2109.273 | 2255.526 | 2690.923 | 1898.855 | 6645.797 | 7431.937 | 5862.028 | 7021.874 | 7341.781 |
| Xanthine | 29234.17 | 42064.29 | 33707.35 | 34544.56 | 40041.68 | 30355.76 | 47405 | 27937.3 | 31482.9 | 52597.1 | 42754.33 | 25280.5 | 26483.01 | 34720.01 | 22636.2 |
| Raltitrexed | 357.9667 | 319.0979 | 244.3234 | 420.8326 | 257.8063 | 376.3727 | 315.6605 | 207.2503 | 346.3097 | 284.9004 | 3350.486 | 2005.584 | 1770.552 | 3886.82 | 2106.182 |
| 3-Nitrotyrosine | 32600.24 | 31322.72 | 30057.83 | 31849.57 | 35271.29 | 31808.11 | 27787.98 | 27880.52 | 33858 | 26247.17 | 30393.04 | 34020.67 | 40901.81 | 36408.72 | 37186.79 |
| Omeprazole | 2113.445 | 3196.929 | 2509.005 | 2339.343 | 3197.596 | 3186.356 | 5628.802 | 3510.566 | 3254.313 | 5437.915 | 4472.64 | 3502.106 | 6090.106 | 5390.154 | 4208.31 |
| Inosinic acid (IMP) | 11957.03 | 9121.822 | 6628.908 | 13029.04 | 8600.747 | 7253.825 | 2876.75 | 6633.03 | 7462.79 | 2621.457 | 7404.676 | 7220.861 | 4846.255 | 7203.025 | 7287.75 |
